# Multi-area activity in mouse motor cortex associated with one- and two-handed oromanual dexterity

**DOI:** 10.64898/2026.04.16.718851

**Authors:** John M. Barrett, Joshua I. Glaser, Andrew Miri, Gordon M. G. Shepherd

## Abstract

Cortical dynamics during goal-directed dexterous hand movements are mainly understood from paradigms involving use of the contralateral hand. To study how movement-related activity changes when the ipsilateral or both hands are used, we exploited a natural form of rodent manual dexterity – food handling – that rodents can perform uni- or bimanually. We sampled kilohertz 3D kinematics as mice used either or both hands to manipulate food, while recording spiking activity in forelimb primary (fl-M1) and secondary (fl-M2) motor cortices, and in a lateral oral and manual (LOM) motor cortex area implicated in oromanual food handling. Unit- and population-level analyses showed that activity in fl-M1 and fl-M2 depended on both laterality (ipsi- vs contralateral) and “manuality” (uni- vs bimanual), with few differences between the two areas. By comparison, activity in LOM was largely laterality- and manuality-invariant. These results demonstrate how activity in multiple areas of mouse motor cortex varies as the same task is performed unimanually with either hand or bimanually with both. Our findings support a model in which fl-M1 and fl-M2 maintain separable information about both forelimbs for bimanual coordination, while LOM encodes ingestion-related forelimb parameters necessary for oromanual coordination.

## INTRODUCTION

Many dexterous behaviors can be performed flexibly with one hand, the other, or both. How is such flexible control manifest in cortical activity? This question has been addressed most extensively in primates, where movement-related cortical activity has been found to be strongly modulated depending on whether the same task is performed with the ipsilateral hand, contralateral hand, or both (Cross et al., 2020; Deo et al., 2024; Donchin et al., 1998, 2002; Ifft et al., 2013; Kermadi et al., 1998, 2000; Rokni et al., 2003; Tanji et al., 1988; Zimnik et al., 2024). In rodents, studies have found bilateral movement encoding during tasks involving bimanual (Jeong et al., 2021) or unimanual (Han et al., 2024; Handa et al., 2024; Rios et al., 2019; Soma et al., 2017, 2019) coordination. Still lacking is an analysis of how spiking activity in multiple areas of rodent cortex is modulated when the same task is performed one-handedly or simultaneously with both hands. This question has also not been studied in the context of complex, ethological forms of dexterity, which may reveal features of cortical population activity not observed in constrained, low-dimensional laboratory tasks (Styr et al., 2026).

Addressing these knowledge gaps requires a natural behavior that rodents can perform unimanually or bimanually, and which they can be trained or induced to switch between the two modes of coordination under experimental control. Of rodents’ natural behavioral repertoire, food handling involves complex movements typically performed in a bimanually coordinated manner (Whishaw et al., 1998), with the thumbs used to hold the smallest food items in a bimanual precision grip (Barrett et al., 2020). However, wild rodents will occasionally handle food unimanually (Missagia et al., 2025). Thus, to exploit this natural behavioral flexibility, we developed a system to gently coax mice to switch between unimanual and bimanual food handling by blocking one hand or the other, or leaving both hands free to access the food.

Food handling engages multiple areas of mouse motor cortex, notably including primary forelimb motor cortex (fl-M1) (Barrett et al., 2022; Xing et al., 2024). Secondary forelimb motor cortex (fl-M2) is also of interest for its circuit connections and functional interactions with fl-M1 (Kristl et al., 2025; Richevaux et al., 2026; Saiki-Ishikawa et al., 2025), as well as differences in unilateral movement encoding between it and fl-M1 (Rios et al., 2019; Soma et al., 2017, 2019). A laterally and anteriorly located area of primary motor cortex, initially characterized as a tongue-jaw area (Mayrhofer et al., 2019; Tamura et al., 2025; Xu et al., 2022), is also involved in food handling behavior (Barrett et al., 2022; Huang et al., 2025; Yang et al., 2023). Optogenetic mapping of this area evokes forelimb as well as orofacial movements (Hira et al., 2015; Khanal et al., 2024; Mayrhofer et al., 2019; Mercer Lindsay et al., 2019). Together, these findings suggest that this area is specialized for behaviors involving coordinated movements of the hand and face. Here, given its roles in forelimb, oral, and particularly food handling actions, we will refer to it as the lateral oral and manual area (LOM).

Thus, to investigate how movement-related activity in fl-M1, fl-M2, and LOM changes as mice handle food unimanually or bimanually, we recorded spiking activity in these areas during food handling while blocking one hand, the other, or neither. Overall, we found that fl-M1 and fl-M2 activity showed partial dependence for ipsilateral versus contralateral and unimanual versus bimanual movements, with few differences between the two areas. In contrast, activity in LOM was largely invariant both to which hand was used, and whether one hand was used or two. This likely reflects different functional roles of these areas, with fl-M1/2 more involved in forelimb movement specification and bimanual coordination, and LOM in oromanual coordination.

## RESULTS

### Head-fixed mice can handle food bimanually or unimanually

A priori, there are four combinatorial possibilities for how activity might vary according to movement laterality - whether it is performed with the ipsilateral or contralateral hand – and “manuality” - whether it is performed with one or two hands (**Figure 1**). The simplest possibility is that activity is the same in all three conditions (ipsilateral, contralateral, or bimanual); i.e., invariant to either movement feature. Activity might depend on which hand is used, but not the number of hands; i.e., laterality dependent. Conversely, activity might depend on the number of hands, but not which hand is used; i.e., manuality dependent. Finally, activity might be different in all three conditions; i.e., show both laterality and manuality dependence.

**Figure 1:**
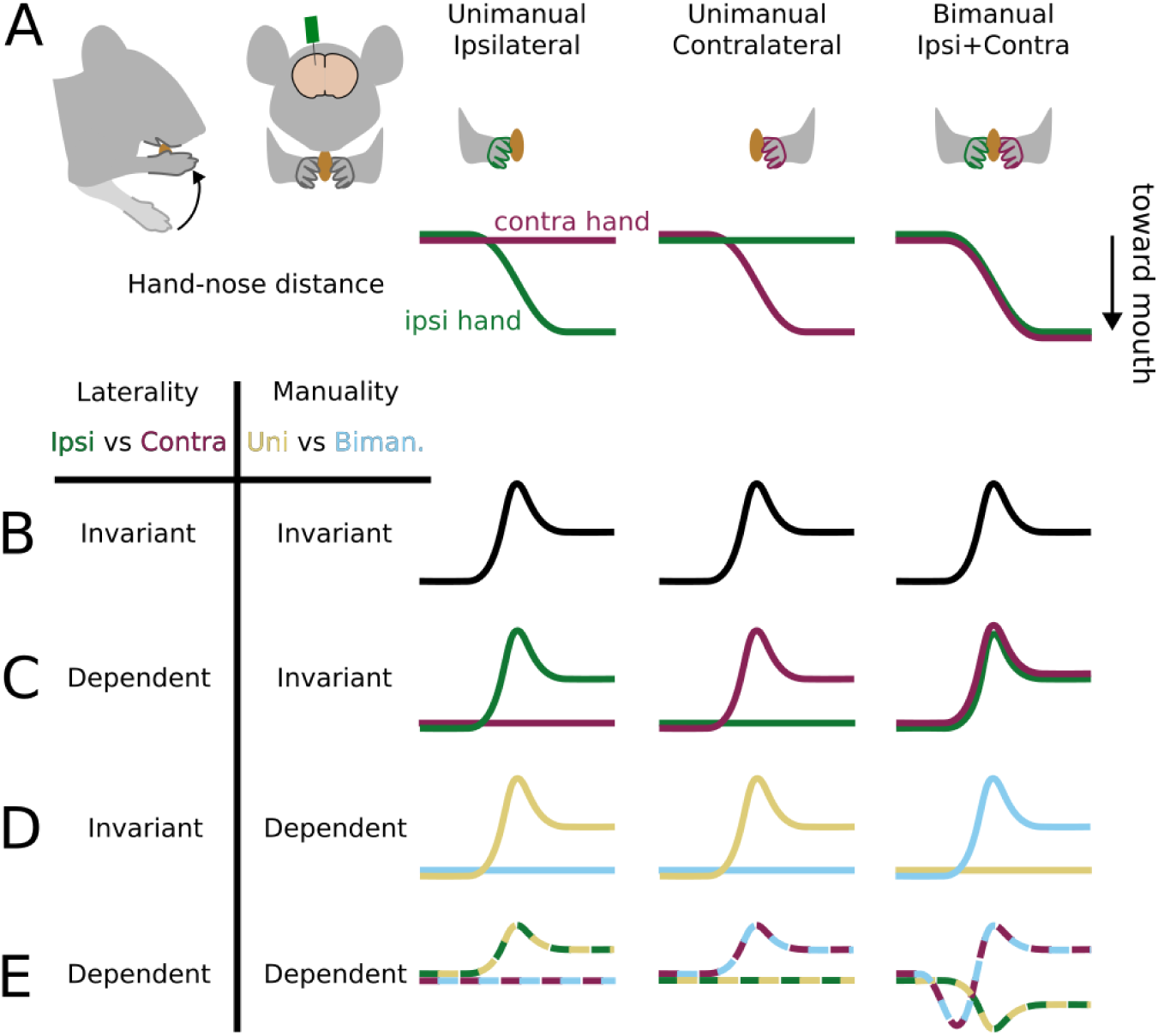
Laterality and manuality dependent and invariant activity. (A) Top row: schematic of cortical recording and food handling conditions. Bottom row: schematic of hand-nose distance for the ipsilateral (green) and contralateral (purple) hands bringing food to the mouth. (B-E) Simplified, non-exhaustive schematic of hypothetical cortical activity patterns under different types of dependence and invariance.

To distinguish these possibilities in the context of head-fixed food handling, we designed a pair of hand blockers to coax mice to use one or the other or both hands for food handling (**Figure 2A**). As previously (Barrett et al., 2022), we used a prism and mirror set-up to stereoscopically film head-fixed mice handling food at kilohertz frame rates. We recorded from one or more cortical areas using 32- or 64-channel linear arrays spanning 1.5 mm of cortical depth, obtaining on average 75 ± 39 (median ± m.a.d., range 10-153) active units (i.e., single units and multiunits) per recording. Most mice, once habituated to the hand blocker setup, successfully performed unimanual food handling with either hand (**Figure 2B,C**). For kinematic analysis, we tracked the nose and third digit on each hand using DeepLabCut (Mathis et al., 2018; Nath et al., 2019) and reconstructed their trajectories in 3D using Anipose (Karashchuk et al., 2021). For each frame, we extracted the distances between each hand and the nose, and between the midpoint of the two hands and the nose.

**Figure 2:**
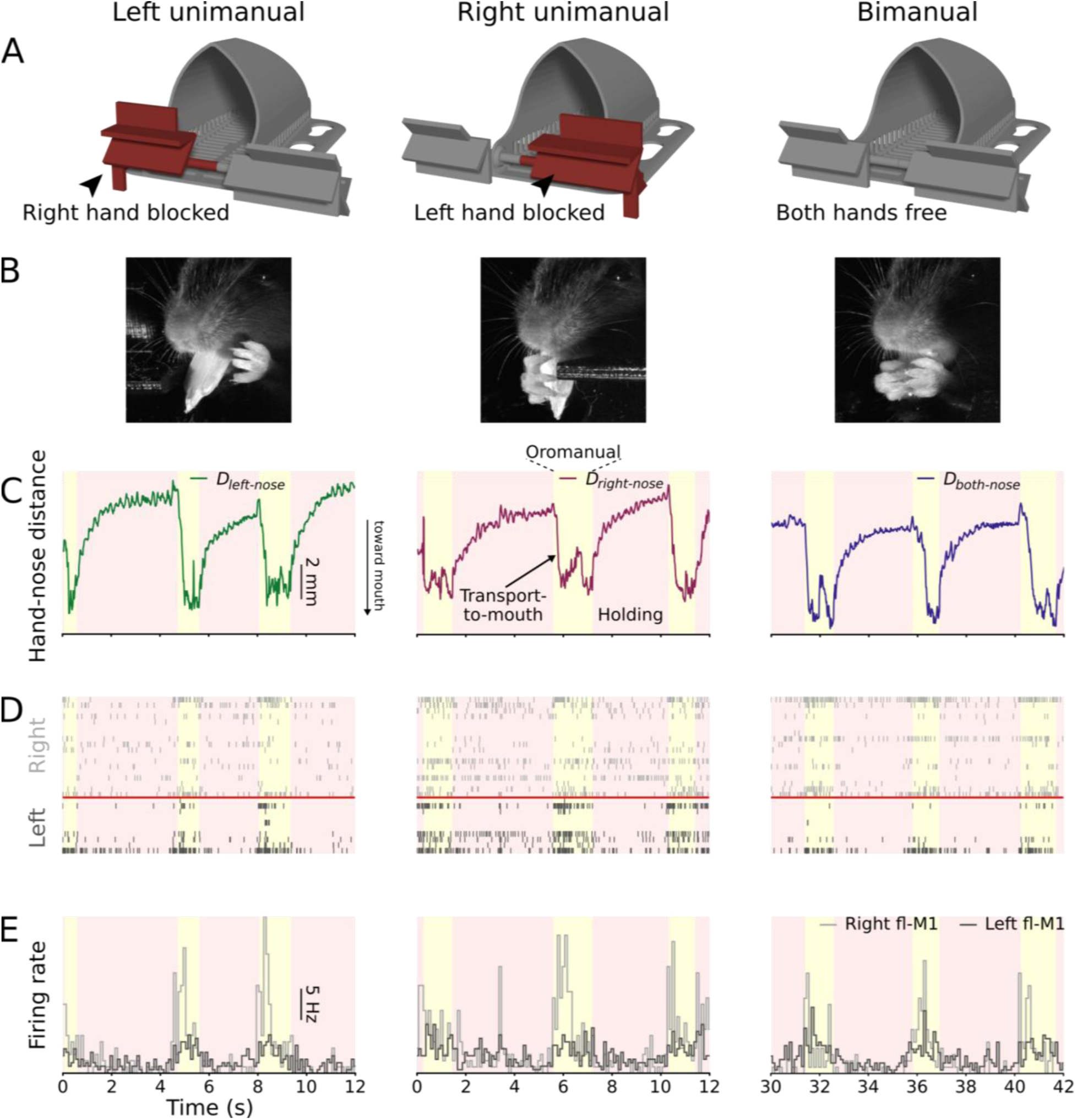
Head-fixed mice can handle food bimanually or unimanually. (A) The hand blocker apparatus consists of a modified arm rest that slides from side to side with flaps that can be rotated and locked in place to prevent one hand or the other from reaching the area immediately in front of and below the mouse’s mouth. (B) Example video frames showing a mouse handling food unimanually with the left hand (left), unimanually with the right hand (middle), and bimanually (right). (C) Example hand-nose distance traces of left unimanual (left, green), right unimanual (middle, purple), and bimanual (right, blue) food handling. (D) Example rasters of all recorded units from a bilateral fl-M1 recording aligned to the kinematic traces in (C). (E) Probe-average firing rate histograms (100 ms bins) of the data in (D).

Rodent food handling involves cycling between two modes: holding/chewing, in which the food is held statically by the hands while the jaw chews, and oromanual/ingestion, in which the hands and jaw work together in a coordinated manner to manipulate food items for ingestion (Barrett et al., 2020, 2024). As shown in the example traces (**Figure 2C**), the holding/chewing and oromanual/ingestion modes can be distinguished by examining the hand-nose distance, as the food is held in closer proximity to the mouth during epochs of oromanual/ingestion than during holding/chewing. Cortical activity tended to increase during oromanual/ingestion and fall during holding/chewing (Barrett et al., 2022) (**Figure 2D,E**).

### Motor cortical activity increases around unimanual and bimanual transport-to-mouth events

To examine cortical activity during unimanual versus bimanual food handling, we first performed event alignment to analyze kinematics and neural activity associated with the “transport-to-mouth” movements occurring at the transition into oromanual/ingestion epochs. These movements, which represent the initiation of periods of active forelimb movements during food handling, are easily detected as sharp decreases in hand-nose distance, and are associated with precisely timed peaks in fl-M1 (Barrett et al., 2022, 2024). During bimanual transports-to-mouth, the hand-nose distance follows a characteristic sigmoidal profile. As shown in the example traces, this is also true for unimanual transports-to-mouth performed with either hand (**Figure 3A,B**). Consistent with prior results, firing rates in fl-M1 increased around bimanual transports-to-mouth. Firing rates increased similarly around both types of unimanual transports-to-mouth (**Figure 3C-E**).

**Figure 3:**
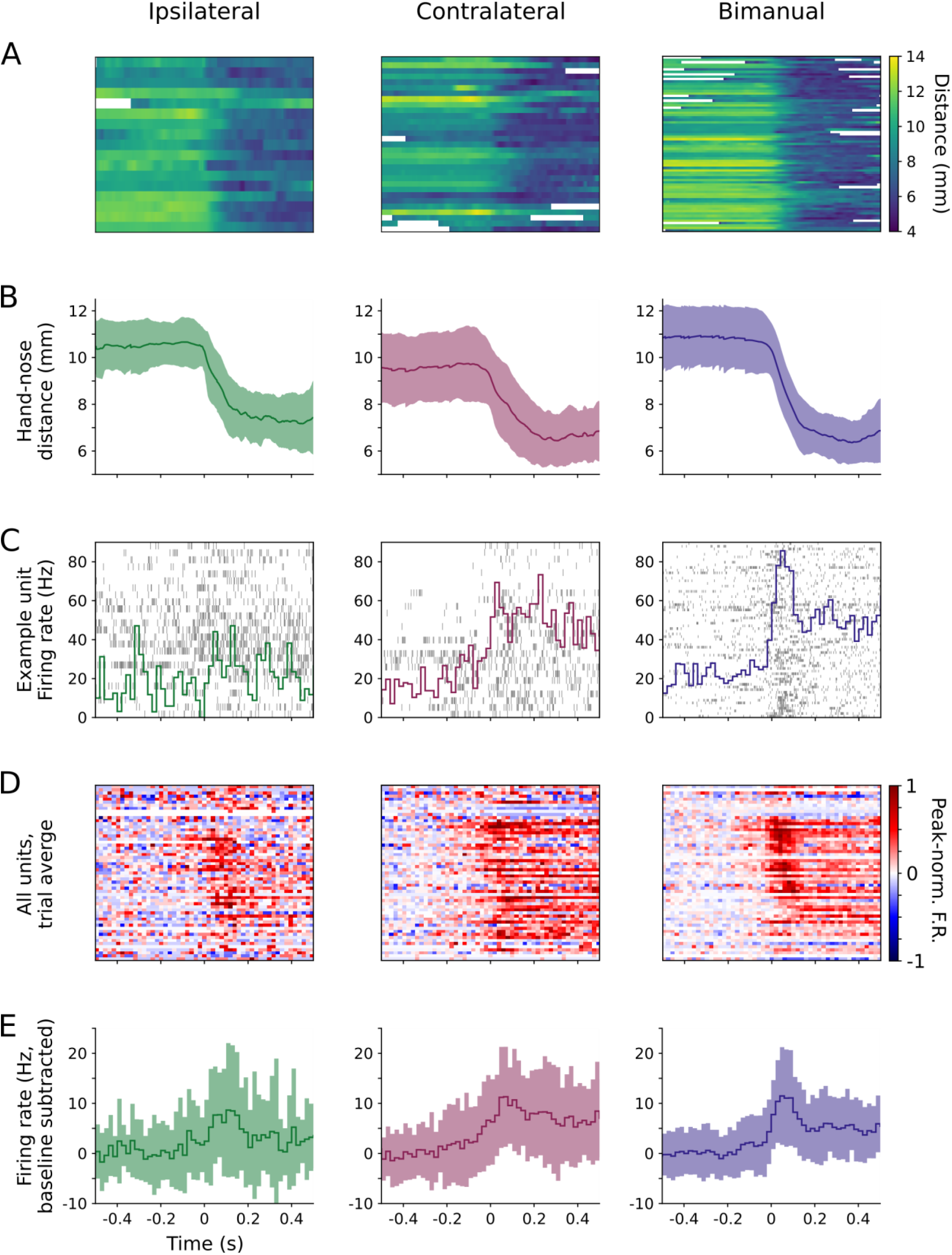
Units in fl-M1 increase in activity around unimanual and bimanual transport-to-mouth events. (A) Hand-nose distance aligned to onsets of ipsilateral (left), contralateral (middle), and bimanual (right) transports-to-mouth, for an example right fl-M1 recording, plotted as a heat map with one row per event. White regions indicate oromanual/ingestion epochs belonging to previous or following holding/chewing-oromanual/ingestion cycles, which were excluded from the aligned traces and PETHs for the purposes of analyzing event-aligned data. (B) Mean (thick line) and S.D. (shaded area) hand-nose distance for the same data in (A). (C) Raster plot (grey lines) and trial-average PETHs (colored lines) for an example fl-M1 unit aligned to the same data in (A-B). (D) Baseline subtracted, peak-normalized, trial-average PETHs for each unit recorded from fl-M1 during the recording associated with (A-C), sorted by time to peak firing around bimanual transports-to-mouth. (E) Grand average (across units and events) baseline-subtracted PETHs for the units in (D).

This pattern held across eight fl-M1 recordings from six mice (**Table 1**, **Figure 4—figure supplement 1**), as grand averages (across probes, days, and mice) showed largely similar activity profiles for all three conditions, and no significant differences in peak firing rates between conditions (**Figure 4A,B,E,F**). Additionally, activity in fl-M2 (four recordings from three mice) showed a broadly similar pattern to fl-M1, also peaking around the onset of oromanual/ingestion (**Figure 4C,E-F**). Finally, LOM (eleven recordings from six mice) displayed relatively delayed and sustained activity during oromanual/ingestion epochs (**Figure 4D-F**). As in fl-M1, probe-average firing rates in fl-M2 and LOM were similar for all three conditions.

**Figure 4:**
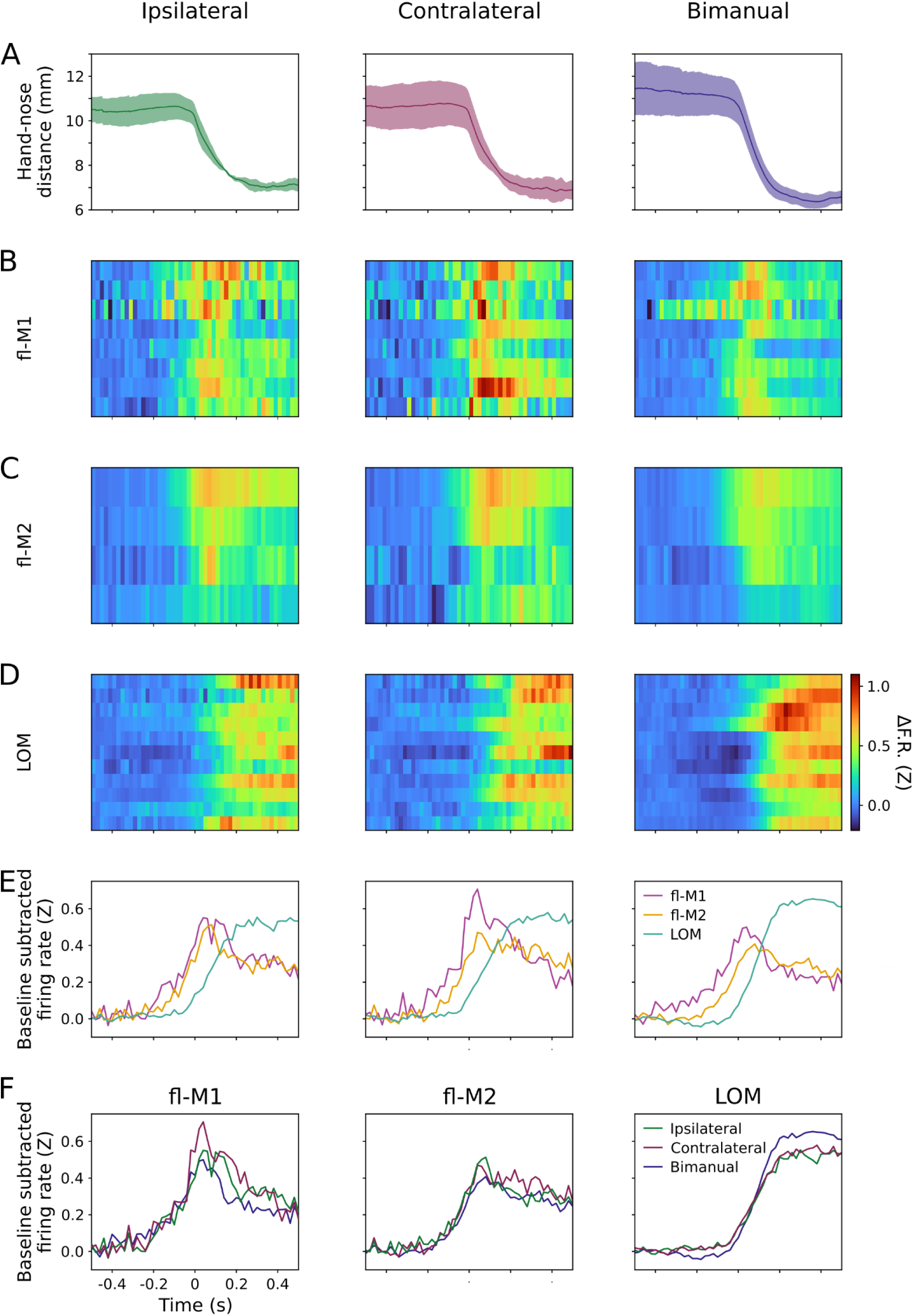
Cortical activity in multiple areas increases around unimanual and bimanual transport-to-mouth events. (A) Mean (lines) and S.D. (shaded area) hand-nose distance traces for 13 recording sessions from 7 mice aligned to ipsilateral (left), contralateral (middle), and bimanual transports-to-mouth (right). (B) Z-scored baseline-subtracted firing rate aligned to transports-to-mouth averaged over events and units for 8 fl-M1 recordings from 6 mice. (C) As (B) for 4 fl-M2 recordings from 3 mice. (D) As (B) for 11 LOM recordings from 6 mice. (E) Z-scored baseline-subtracted firing rate aligned to transports-to-mouth averaged over probes, days, then mice in fl-M1 (violet), fl-M2 (gold), and LOM (teal). (F) As (E), but arranged by area and colored by condition. Left: fl-M1, middle: fl-M2, right: LOM. Green: ipsilateral, purple: contralateral, blue: bimanual.

**Table 1:** mice appearing in this paper.

| # | Age | Weight | Sex | Analyses |  |  | PCA Conditions |
| --- | --- | --- | --- | --- | --- | --- | --- |
|  |  |  |  | fl-M1 | fl-M2 | tj-M1 |  |
| 1 | 231 | 23.0 | F |  |  | PCA | Ipsi↔Bi |
| 2 | 206 | 26.8 | M | PCA |  |  | Ipsi↔Contra |
| 3 | 140 | 18.7 | F | PETH, PCA, Corr, GLM | PCA, Corr, GLM | PETH, PCA, Corr, GLM | Contra↔Bi, Ipsi↔Bi, Ipsi↔Contra |
| 4 | 117 | 20.6 | M | PETH, Corr, GLM | PETH, PCA, Corr, GLM |  | Contra↔Bi, Ipsi↔Bi, Ipsi↔Contra |
| 5 | 118 | 25.6 | M |  |  | PETH, PCA, Corr, GLM | Ipsi↔Bi |
| 6 | 109 | 19.5 | F | PETH, Corr, GLM |  | PETH, PCA, Corr, GLM | Contra↔Bi, Ipsi↔Bi |
| 7 | 116 | 18.9 | F |  | PCA |  | Contra↔Bi, Ipsi↔Bi |
| 8 | 153 | 24.4 | M | PETH, PCA, Corr, GLM | PETH, Corr, GLM | PETH, Corr, GLM | Contra↔Bi |
| 9 | 150 | 25.3 | M | PETH, Corr, GLM |  | PETH, PCA, Corr, GLM | Contra↔Bi, Ipsi↔Bi, Ipsi↔Contra |
| 10 | 150 | 24.0 | M | PETH, Corr, GLM | PETH, Corr, GLM | PETH, Corr, GLM |  |

These overall average activity profiles in the three areas and three conditions show that activity was generally not strongly lateralized during unimanual food handling, but instead was largely similar for ipsilateral unimanual, contralateral unimanual, and bimanual handling. The overall activity profiles in fl-M1 and LOM resembled those found previously for these areas (Barrett et al., 2022). The activity profile in fl-M2 resembled fl-M1 more than LOM (**Figure 4E**).

### Activity is more laterality- and manuality-dependent in forelimb areas than in LOM

Averaging across all units in a given recording can obscure differences in individual unit responses, in which some units might respond selectively during ipsilateral, contralateral, or bimanual movements (**Figure 3D**). Hence, for each unit, we assessed whether its firing rate significantly increased around transports-to-mouth using a bootstrap test (**Methods**). This showed that some units showed significant increases in firing around transports-to-mouth in only one condition (ipsilateral, contralateral, or bimanual), in two out of three conditions, or in all three conditions (**Figure 5A-C**). In fl-M1, the largest group of units significantly increased firing during contralateral transports-to-mouth only (22.8%). Bimanual only was the next largest (20.7%), then significant during all (19.9%), then significant during bimanual and contralateral transports-to-mouth (14.9%; **Figure 5D-F**). Units with significant firing rate changes during ipsilateral and bimanual but not contralateral was the rarest category, consistent with largely contralaterally projecting outputs from fl-M1. The overall pattern was similar in fl-M2, but with relatively fewer bimanual only units. LOM showed a very different pattern, with the largest single category being units significant during all types of transports-to-mouth, and roughly even proportions of the remaining categories. The difference in proportions between areas was significant (*χ*^2^ test of independence: *χ*^2^ = 31.3, *d.f.* = 12, *p* = 0.002).

**Figure 5:**
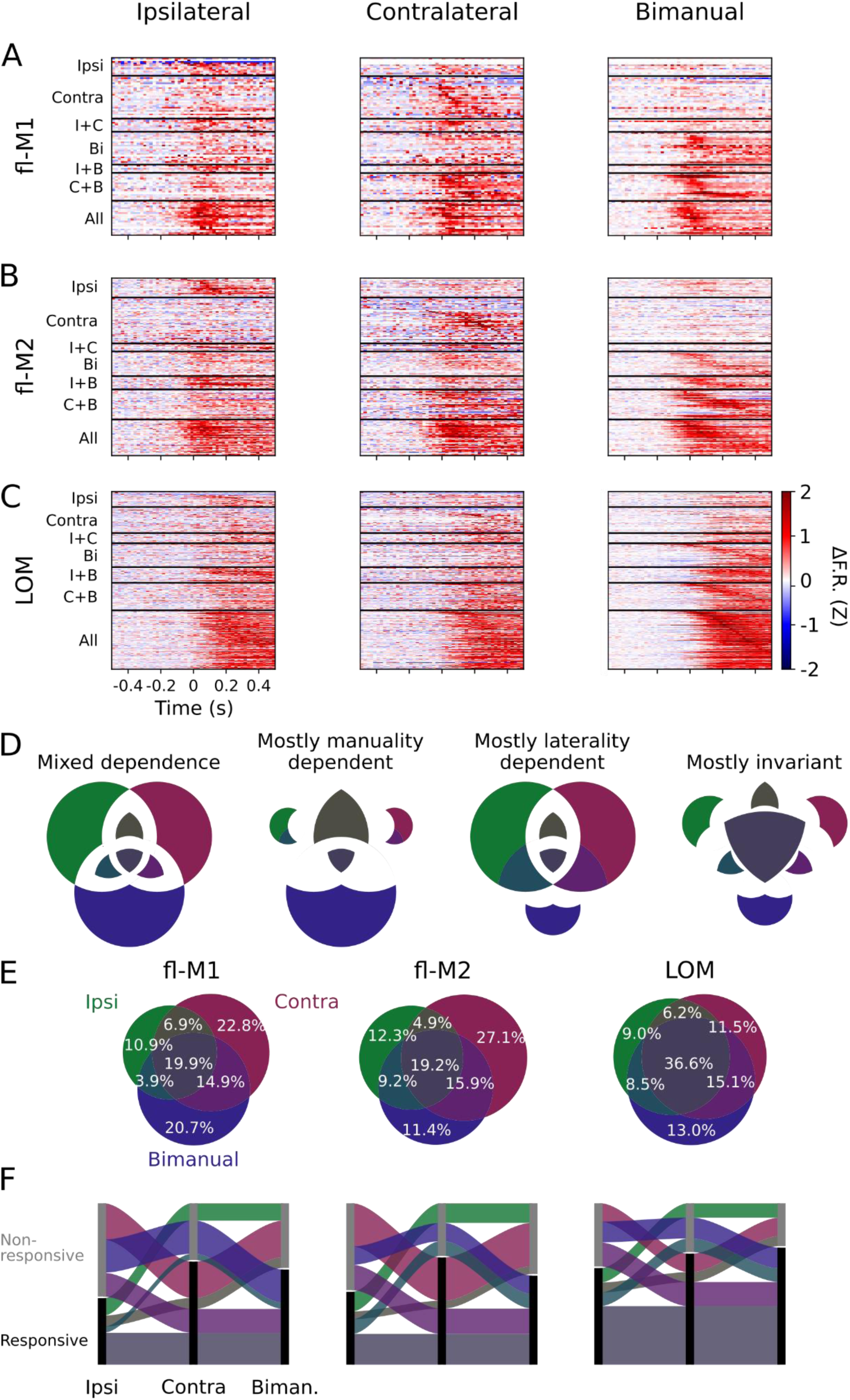
LOM shows a higher proportion of non-selective significant responses compared to forelimb areas. (A) Trial-average, baseline-subtracted, Z-scored PETHs for all recorded fl-M1 units (pooled across recordings and mice) aligned to ipsilateral (left), contralateral (middle), and bimanual (right) transports-to-mouth. Units are organized by condition(s) for which they show significant firing rate increases. (B) As (A), but for fl-M2 (C) As (A), but for LOM (D) Exploded Venn diagrams showing predicted proportions of significant responses under different types of dependence (qualitatively, and assuming symmetric laterality preferences). (E) Venn diagrams showing proportions of units (calculated per recording, then averaged over probes, days, and mice) that are significantly responsive during only one type of transport-to-mouth, to two types, or to all three in fl-M1 (left, 8 recordings from 6 mice), fl-M2 (middle, 4 recordings from 3 mice), and LOM (right, 11 recordings from 6 mice). (F) The same data as in (E) re-plotted as a Sankey diagram. Note the relative size of the dark red (contralateral only) band in fl-M1/fl-M2 and grey (responsive during all) band in LOM.

The above analysis considers responses during each condition in isolation. As a complementary approach to directly compare responses in each condition, for each unit responsive to any type of transport-to-mouth we calculated a preference index, (*A***-***B*)/(*A*+*B*), where *A* is the unit’s average firing rate during a 100 ms window surrounding its time of peak firing during condition A, and *B* defined similarly for condition B. We used this metric to gauge both laterality preference, by comparing responses during ipsilateral versus contralateral transports-to-mouth, and manuality preference, by comparing responses during bimanual transports-to-mouth to each unit’s preferred unimanual condition (**Figure 6A**). The distribution of ipsilateral versus contralateral preferences significantly differed between fl-M2 and LOM (permutation test on Kolmogorov-Smirnov distance, *p* = 0.04, **Figure 6B**), with fl-M2 having more units with strong ipsilateral or contralateral preferences. There were no significant differences between fl-M1 and fl-M2 (*p* = 0.65) or LOM (*p* = 0.39). This held when comparing the distributions of the absolute values of ipsilateral versus contralateral preference indices, which significantly differed between fl-M2 and LOM (*p* < 0.0001), with larger laterality preference indices in the former and smaller laterality preference indices in the latter (**Figure 6C**). Finally, unimanual versus bimanual preference distributions were significantly more skewed towards unimanual preferences in fl-M1 and fl-M2 and toward bimanual preferences in LOM, with no difference between fl-M1 and fl-M2 (fl-M1 vs fl-M2: *p* = 0.86; fl-M1 vs LOM: *p* = 0.002; fl-M2 vs LOM: *p* < 0.0001; **Figure 6D**).

**Figure 6:**
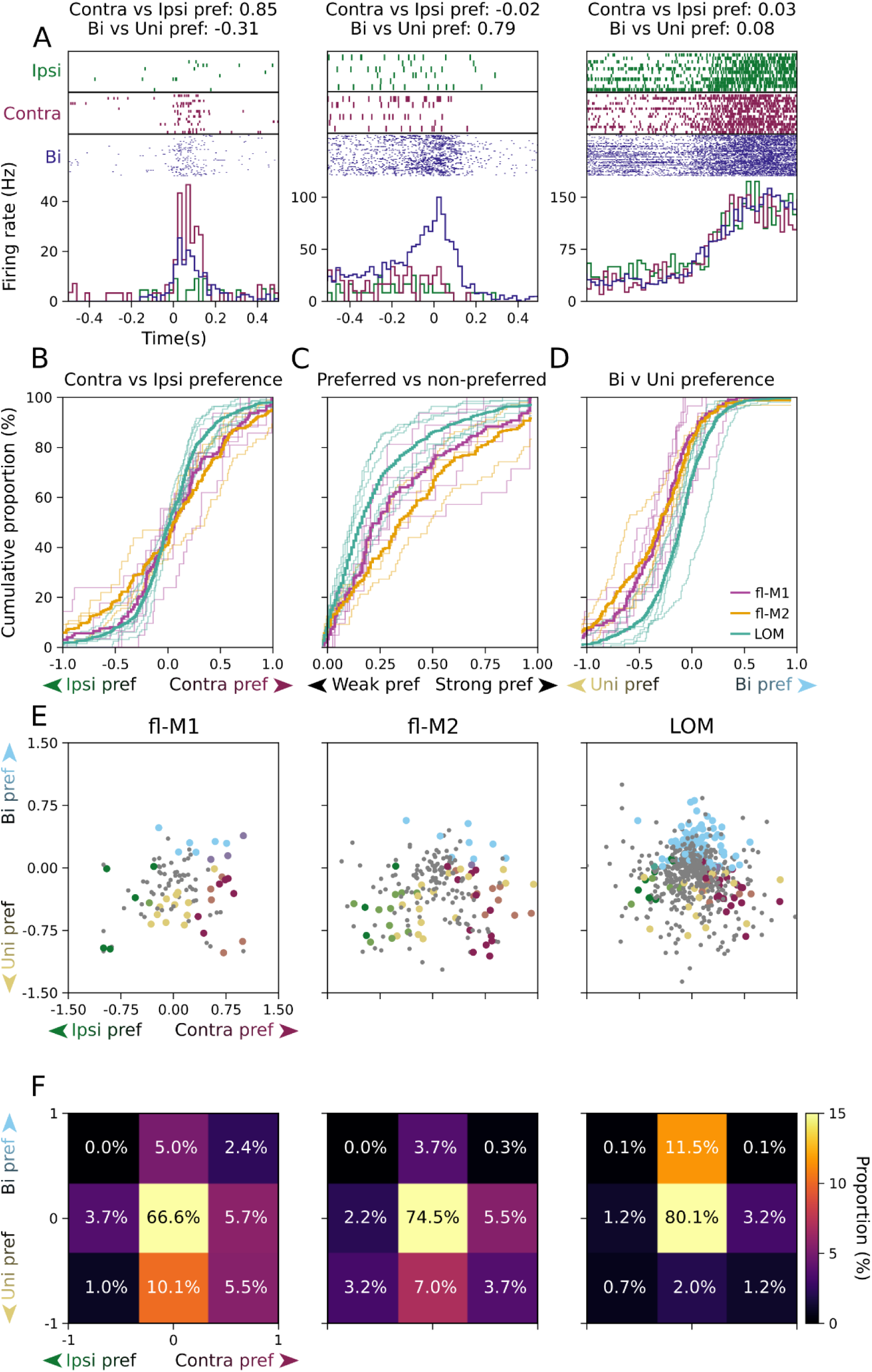
The fl-M2 shows the most lateralized transition-aligned activity, LOM the least lateralized. (A) example raster plots and PETHs (green: ipsilateral, purple: contralateral, blue: bimanual) for three example units - one with a strong contralateral preference but a weak bimanual preference (left), one with a weak ipsilateral preference but a strong bimanual preference (middle), and one with no strong preferences (right). (B) Cumulative distributions of preference indices for contralateral versus ipsilateral transports-to-mouth for fl-M1 (violet), fl-M2 (gold), and LOM (teal). (C) As (B), for absolute value of contralateral versus ipsilateral preference indices. (D) As (B), for bilateral transports-to-mouth versus each unit’s preferred unimanual transports-to-mouth. (E) Scatter plot of laterality preference indices vs manuality preference indices for all recorded units in fl-M1 (left, 8 recordings from 6 mice), fl-M2 (middle, 4 recordings from 3 mice), and LOM (right, 11 recordings from 6 mice). Small grey dots indicate no significant preference. Large colored dots indicate significant ipsilateral preference (green), ipsilateral and unimanual preference (lime), unimanual preference (yellow), contralateral and unimanual preference (peach), contralateral preference (purple), contralateral and bimanual preference (lavender), bimanual preference (light blue), or ipsilateral and bimanual preference (turquoise). (F) 2D histogram of the data in (E).

To examine the joint distribution of laterality and manuality preferences, we categorized each unit where we could detect significant differences between its responses during ipsilateral versus contralateral (or bimanual versus unimanual) transports-to-mouth (Mann-Whitney *U* test, *p* < 0.05, uncorrected) as having a strong laterality (or manuality) preference (**Figure 6E-F**). In all three areas, we could not detect strong preferences in the majority of units (67% in fl-M1, 75% in fl-M2, and 80% in LOM), but where we did find strong preferences the distributions were skewed more towards unimanual and/or contralateral preferring units in fl-M1 and fl-M2, and towards bimanual preferring units with no laterality preference in LOM. The histograms of these categories (**Figure 6F**) were significantly different in fl-M1 and fl-M2 compared to LOM (permutation test on earth mover’s distance, fl-M1 vs LOM: *p* = 0.01, fl-M2 vs LOM: *p* = 0.009), but not between fl-M1 and fl-M2 (*p* = 0.71). Altogether, these analyses of responsiveness and preference index show greater variation in firing with laterality in fl-M1 and especially fl-M2, and less in LOM. Units in fl-M1/2 were more biased towards unimanual responses, particularly contralateral unimanual responses, whereas LOM units were slightly biased towards bimanual responses.

### Population dynamics are more conserved across conditions in LOM than forelimb motor cortices

Individual unit responses give only limited insight into how neurons work together as a population to encode movement parameters. To identify major features of the population response, we applied principal components analysis (PCA) to the transport-to-mouth aligned activity. PCA was performed on event-aligned average responses for units from each region, separately during ipsilateral, contralateral, and bimanual transports-to-mouth. This analysis was restricted to recordings with at least 40 identified units and at least two conditions with 15 or more transports-to-mouth each (**Methods, Table 1**). **Figure 7A** shows the top three principal components (PC) over time for the bimanual condition in an example fl-M1 recording, and the top two PCs plotted against each other. The top PC in particular resembles the probe-average activity, transiently peaking around transport-to-mouth, and the neural activity follows a rotational trajectory through the space spanned by the top two PCs. Similarly, for LOM (**Figure 7B**), the top PC shows sustained increases during oromanual events and rotational dynamics are also observed in the top two PCs. Overall, the top PC contributed a large fraction of the explained variance, but the amount significantly differed depending on condition and area (linear mixed effects model, **Figure 7C,D**, **Table 2**). In particular, the top PC accounted for a greater portion of the explained variance in LOM (67%, average across conditions) compared to fl-M1 (42%, *p* = 0.005) or fl-M2 (45%, *p* = 8.9 × 10^−6^). This suggests lower dimensional activity during transports-to-mouth in LOM, and accordingly there was a significant effect of area on the participation ratio (a measure of dimensionality, Gao et al., 2017), which was lower in LOM than fl-M1 (*p* = 0.003) or fl-M2 (*p* = 7.5 × 10^−8^).

**Figure 7:**
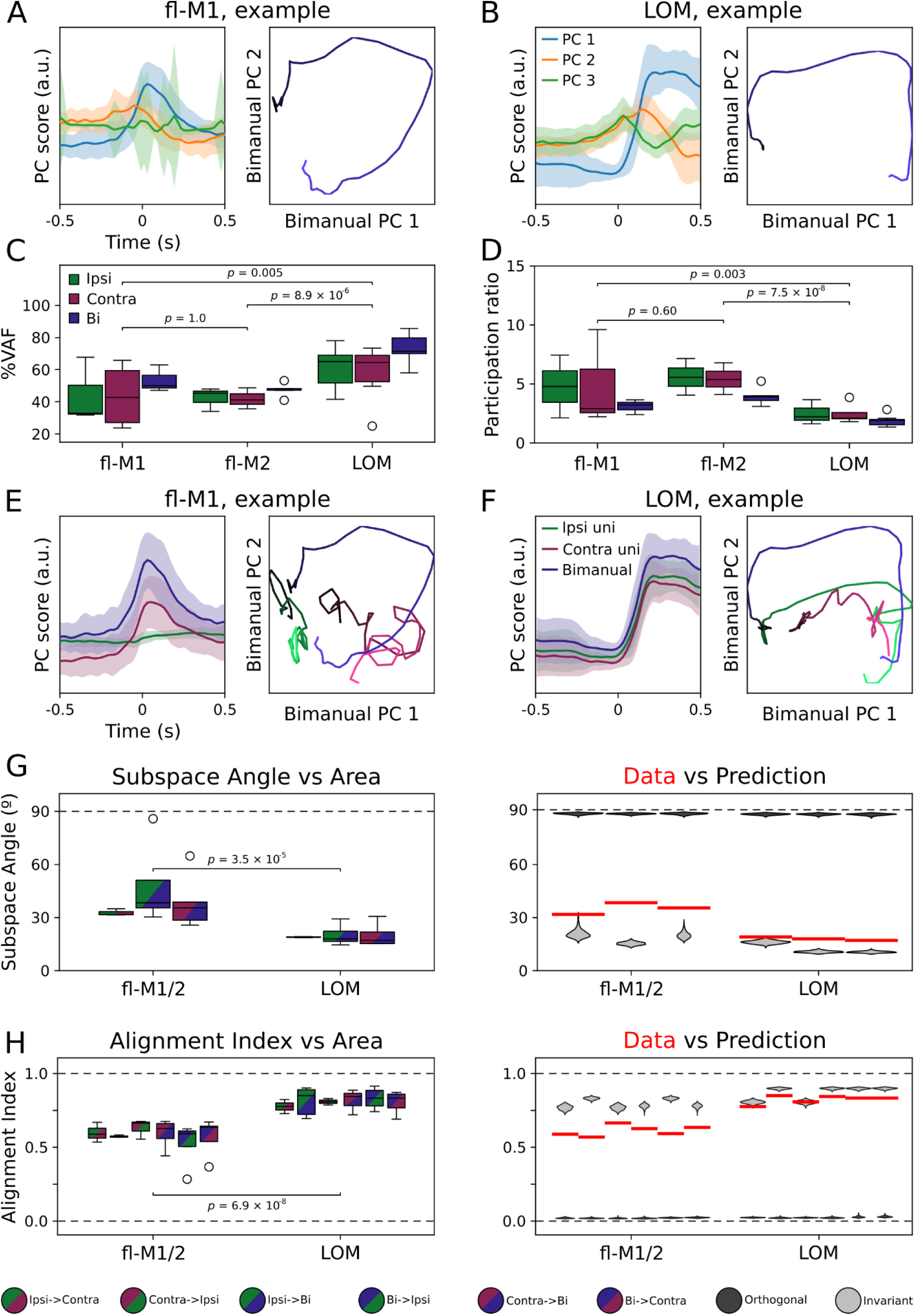
Population dynamics are more conserved across conditions in LOM than forelimb motor cortices. (A) Mean bimanual transport-to-mouth aligned activity for an example fl-M1 recording projected onto the top three principal components recovered from the data (left), and the activity along the top two principal components plotted against each other over time (right) for transports-to-mouth (blue). (B) As (A) for an example LOM recording. (C) Boxplot of fraction of explained variance accounted for by the top principal component of transport-to-mouth aligned average activity for all mice (see **Table 1** for mice included for each condition), organized by area then condition. (D) As (C), but organized by condition then area. (E) Left: Event aligned neural activity associated with bimanual transports-to-mouth in an example fl-M1 recording projected onto the top principal component identified from ipsilateral (green), contralateral (purple), and bimanual (blue) transport-to-mouth responses. Right: Event aligned neural activity associated with ipsilateral (green), contralateral (purple), and bimanual (blue) transports-to-mouth projected onto the top two bimanual principal components. Blue traces are the same as in (A). (F) As (E) for an example LOM recording. (G) Left: Boxplot of subspace angles between the top principal components for each area and condition pair. Right: Expected distributions of subspace angles under laterality/manuality invariance (light gray) and maximum laterality/manuality dependence (i.e. orthogonality, dark gray) compared to the real data (red bars: condition pair/area medians from left panel). Areas fl-M1 and fl-M2 have been pooled to ensure at least two mice were included for each condition pair. (H) As (G) for alignment index between the top ten principal components.

**Table 2:** statistical analyses (linear mixed-effects models)

| Top principal component variance vs area and condition |  |  |  |  |  |
| --- | --- | --- | --- | --- | --- |
|  | SS | MS | df | F | p |
| **Region** | 0.20 | 0.10 | 2 | 9.7 | *0.0003* |
| **Condition** | 0.12 | 0.06 | 2 | 6.0 | *0.006* |
| **Interaction** | 0.02 | 0.01 | 4 | 0.49 | 0.74 |

| Participation ratio vs area and condition |  |  |  |  |  |
| --- | --- | --- | --- | --- | --- |
|  | SS | MS | df | F | p |
| Region | 66.4 | 33.2 | 2 | 15.3 | *0.0002* |
| Condition | 13.5 | 6.7 | 2 | 3.1 | 0.06 |
| Interaction | 4.1 | 1.0 | 4 | 0.47 | 0.75 |
| Subspace angle vs area (fl-M1, fl-M2, or LOM) and condition pair |  |  |  |  |  |
|  | SS | MS | df | F | p |
| Area | 1668.6 | 834.3 | 2 | 16.36 | $8.4 \times 10^{-5}$ |
| Condition Pair | 470.6 | 235.3 | 2 | 4.61 | *0.03* |
| Interaction | 540.23 | 135.1 | 4 | 2.65 | 0.08 |
| Subspace angle vs region (fl-M1/2 or LOM) and condition pair |  |  |  |  |  |
|  | SS | MS | df | F | p |
| Region | 1763.6 | 1763.6 | 1 | 25.85 | $3.1 \times 10^{-5}$ |
| Condition Pair | 198.56 | 99.28 | 2 | 1.46 | 0.25 |
| Interaction | 148.74 | 74.37 | 1 | 1.09 | 0.08 |
| Alignment index vs area and condition pair |  |  |  |  |  |
|  | SS | MS | df | F | p |
| Area | 0.23 | 0.12 | 2 | 30.63 | $1.3 \times 10^{-8}$ |
| **Condition Pair** | 0.07 | 0.01 | 5 | 3.76 | *0.008* |
| **Interaction** | 0.04 | 0.004 | 10 | 0.96 | 0.49 |
| **Alignment index vs region and condition pair** |  |  |  |  |  |
|  | **SS** | **MS** | **df** | **F** | **p** |
| **Region** | 0.22 | 0.22 | 1 | 40.57 | $6.9 \times 10^{-8}$ |
| **Condition Pair** | 0.04 | 0.008 | 5 | 1.40 | 0.25 |
| **Interaction** | 0.01 | 0.002 | 5 | 0.37 | 0.87 |
| **Population activity structure correlation vs area and condition pair** |  |  |  |  |  |
|  | **SS** | **MS** | **df** | **F** | **p** |
| **Area** | 0.79 | 0.40 | 2 | 87.47 | $1.1 \times 10^{-18}$ |
| **Pair** | 0.03 | 0.02 | 2 | 3.76 | *0.03* |
| **Interaction** | 0.03 | 0.008 | 4 | 1.70 | 1.6 |
| **Decoding accuracy vs area, dimension (X, Y, or Z), laterality, and manuality** |  |  |  |  |  |
|  | **SS** | **MS** | **df** | **F** | **p** |
| **Area** | 1.74 | 0.87 | 2 | 32.78 | $4.0 \times 10^{-12}$ |
| **Dimension** | 6.54 | 3.27 | 2 | 123.32 | $5.4 \times 10^{-39}$ |
| **Laterality** | 0.007 | 0.007 | 1 | 0.23 | 0.62 |
| **Manuality** | 0.30 | 0.30 | 1 | 11.45 | *0.0008* |

| Percent change in decoding accuracy vs laterality and generalization type (cross-body or cross-condition) |  |  |  |  |  |
| --- | --- | --- | --- | --- | --- |
|  | SS | MS | df | F | p |
| Area | 1.67 | 0.83 | 2 | 3.53 | *0.03* |
| Laterality | 0.14 | 0.14 | 1 | 0.58 | 0.45 |
| Generalization type | 0.21 | 0.21 | 1 | 0.88 | 0.35 |

Next, we assessed how consistent the population activity structure was across conditions by testing how well the PCA projections generalized across conditions. **Figure 7E** shows activity from bimanual transports-to-mouth projected onto the top PC from the ipsilateral, contralateral, and bimanual conditions for the same example recording (left traces), as well as the activity from each condition projected onto the top two PCs from the bimanual condition (right X-Y plot). Note how the same activity results in reduced magnitude traces depending on which PC space it is projected into and, conversely, unimanual activity projected into the bimanual PC space follows very different trajectories to the bimanual activity. By contrast, in LOM (**Figure 7F**), the bimanual activity results in almost the same trace regardless of which PC space it is projected into and the unimanual activity proceeds in roughly the same rotational direction within the space spanned by the top two bimanual PCs. This suggests that the PCs are more similar across conditions for LOM, but more variable across conditions for fl-M1 and fl-M2.

To see if this pattern held in general, we quantified, for each pair of conditions (ipsilateral vs contralateral, ipsilateral vs bimanual, and contralateral vs bimanual), the first principal angle between the subspaces spanned by the top PCs (**Figure 7G**), and the alignment index for the top 10 PCs (**Figure 7H**, **Methods**, (Elsayed et al., 2016)). If activity is laterality invariant, we would expect the subspace angles to be close to zero and the alignment index to be close to one when comparing ipsi- to contralateral subspaces, and likewise when comparing either unimanual subspace to the bimanual subspace if activity is manuality invariant. Due to the stricter inclusion criteria, some condition pairs had only mouse included in fl-M1 or fl-M2 when these areas were analyzed separately, hence these areas are pooled in **Figure 7**. The results held if all three areas were analyzed separately, and there were no differences between fl-M1 and fl-M2 (**Figure 7—figure supplement 1**). Subspace angles were lower in LOM than fl-M1/2 (linear mixed effects model; *p* = 3.1 × 10^−5^; **Table 2**), and alignment indices higher (*p* = 6.9 × 10^−8^). There was no significant main effect of condition pair (subspace angle: *p* = 0.26, alignment index: *p* = 0.25), indicating similar degrees of alignment between ipsilateral and contralateral subspaces as between either unimanual subspace and the bimanual subspace. This is consistent with manuality dependence not simply being driven by laterality dependence, as in that case one would expect to find greater alignment between the bimanual subspace and one of the unimanual subspaces compared to the other.

To assess how well the observed data matched predictions for laterality/manuality invariant or dependent activity, for each condition pair in each area, we bootstrapped estimated distributions of the expected subspace angles and alignment indices under the assumption of perfectly aligned or perfectly orthogonal subspaces (**Figure 7G-H, Methods**). Averaging across conditions, the subspace angles and alignment indices were closer to the invariant prediction in LOM than fl-M1/M2 (permutation test: subspace angle, *p* = 0.003; alignment index: *p* < 0.0001). In summary, these analyses show greater overlap between conditions in the LOM activity subspaces than fl-M1 and fl-M2 activity subspaces. This is consistent with greater laterality and manuality dependence in fl-M1 and fl-M2 compared to LOM.

### Population correlation structure reorganizes between conditions in forelimb but not LOM

The above analyses only considered trial-average activity aligned to transports-to-mouth. As a complementary approach to assess how population activity varies on a moment-to-moment basis during all phases of food handling, we calculated the correlation coefficients between each pair of units throughout continuous bouts of ipsilateral, contralateral, and bimanual food handling (**Figure 8**). In an example fl-M1 recording, there are clear differences in the structure of the pairwise correlation matrices between conditions (**Figure 8A**), but these are more subtle for simultaneously recorded activity in LOM (**Figure 8B**). To quantify this, we calculated the correlation coefficient between the pairwise correlation matrices for each condition (**Figure 8C**). Overall, there were significant main effects of area and of condition pair on similarity of population correlation matrices between conditions (linear mixed-effects model, **Table 2**, **Figure 8D**), with follow-up tests showing higher between-condition correlation in LOM than fl-M1 (*p* = 2.6 × 10^−14^) or fl-M2 (*p* = 9.2 × 10^−20^), but not between fl-M1 and fl-M2 (*p* = 0.33). Averaging across condition pairs, the between-condition correlation coefficient in LOM was 0.89, indicating almost entirely preserved population correlation structure. It was 0.63 in fl-M1 and 0.64 in fl-M2, indicating some degree of shared activity structure. Thus, population activity during food handling partially reorganizes in forelimb areas according to which hands are involved, whereas it largely retains its structure in LOM.

**Figure 8:**
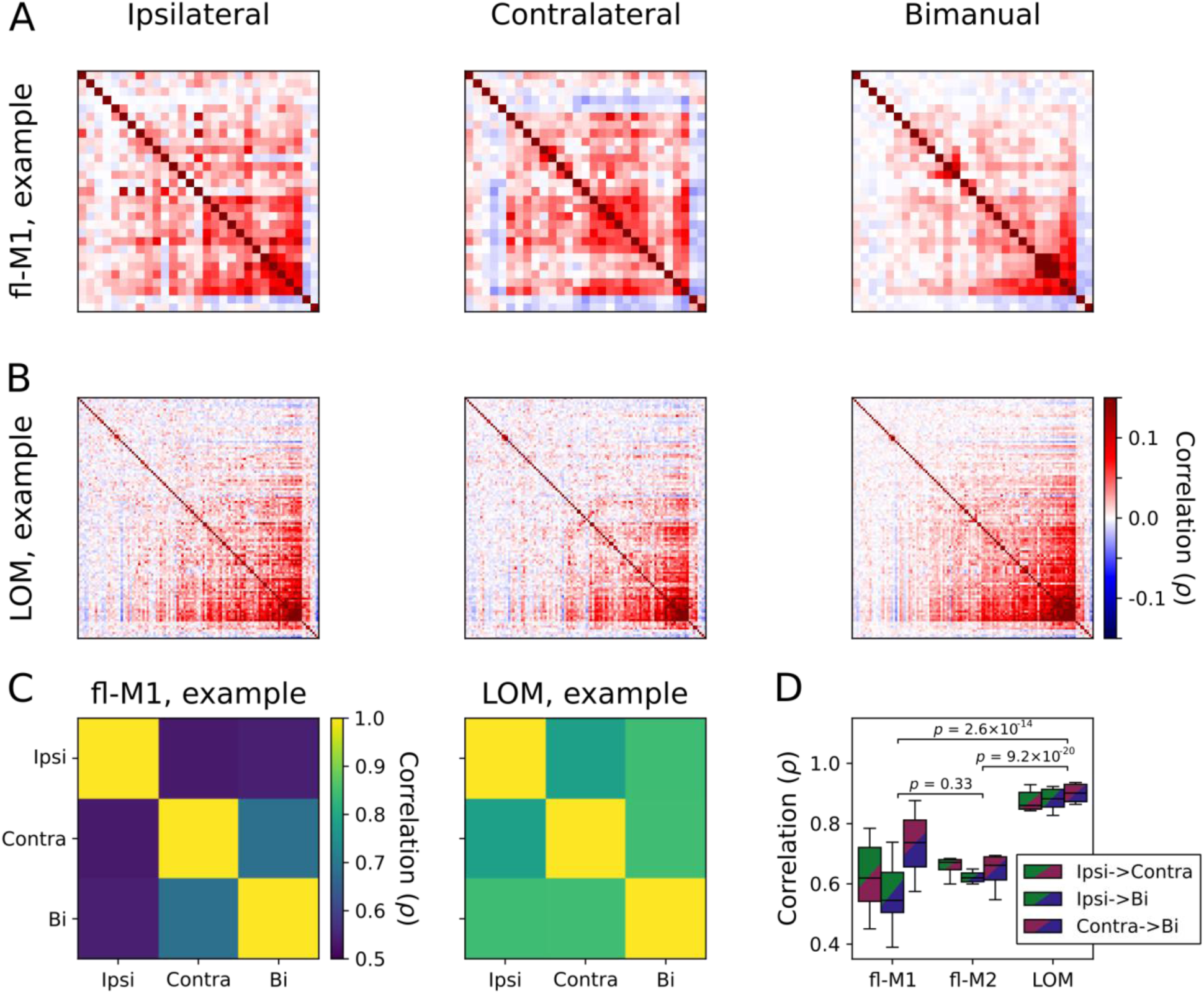
Population activity throughout food handling reorganizes between conditions in forelimb motor cortices but not LOM. (A) Pairwise correlations for an example fl-M1 recording during continuous ipsilateral (left), contralateral (middle), and bimanual (right) food handling. Ordering of units is consistent between panels. (B) As (A) but for simultaneously recorded LOM activity. (C) Correlation between the fl-M1 (left) and LOM (right) population correlation matrices for each condition, for the recording in (A-B)= (D) Boxplot of correlation between population correlation matrices for each pair of conditions for all recordings (fl-M1: 7 recordings from 6 mice, fl-M2: 6 recordings from 4 mice; LOM: 11 recordings from 6 mice), organized by area.

### Kinematic decoding is preserved between conditions in LOM but not forelimb motor cortices

To understand how these changes in population activity structure affect how movements can be read out from neural activity, we decoded continuously recorded forelimb kinematics from neural activity throughout all phases of food handling using general linear models (GLMs) fit to lagged windows (equivalent to Wiener filters (Carmena et al., 2003; Glaser et al., 2020)), as previously (Barrett et al, 2022). In each condition (ipsilateral, contralateral, and bimanual) and for each limb, we fit a GLM that decoded the 3D position of the hands throughout food handling (including holding/chewing and oromanual/ingestion, but excluding non-food handling behavior) from neural activity in each area (**Figure 9A, Methods**). As expected, kinematics could be decoded above chance from all three areas (**Figure 9B**). To quantify decoding accuracy, we calculated the cross-validated *R*^2^ for held-out data. Decoding accuracy varied significantly across cortical areas, decoded dimensions, and between unimanual and bimanual conditions, but was not significantly different for the two hands (linear mixed-effects model, **Figure 9C**, **Table 2**). Follow-up tests showed that decoding was significantly better in LOM compared to fl-M1 (*p* = 6.5 × 10^−8^) and fl-M2 (*p* = 0.04), and better in fl-M2 than fl-M1 (*p* = 0.008). Decoding was better for the Y (up-down) dimension than Z (rostral-caudal), which in turn was better than X (left-right; X vs Y: *p* = 2.6 × 10^−30^, X vs Z: *p* = 0.002, Y vs Z: *p* = 8.4 × 10^−17^). This is likely due to the largest and most consistent movements (transports-to-mouth and lowering from mouth) involving mostly vertical translation, with a small rostrocaudal component.

**Figure 9:**
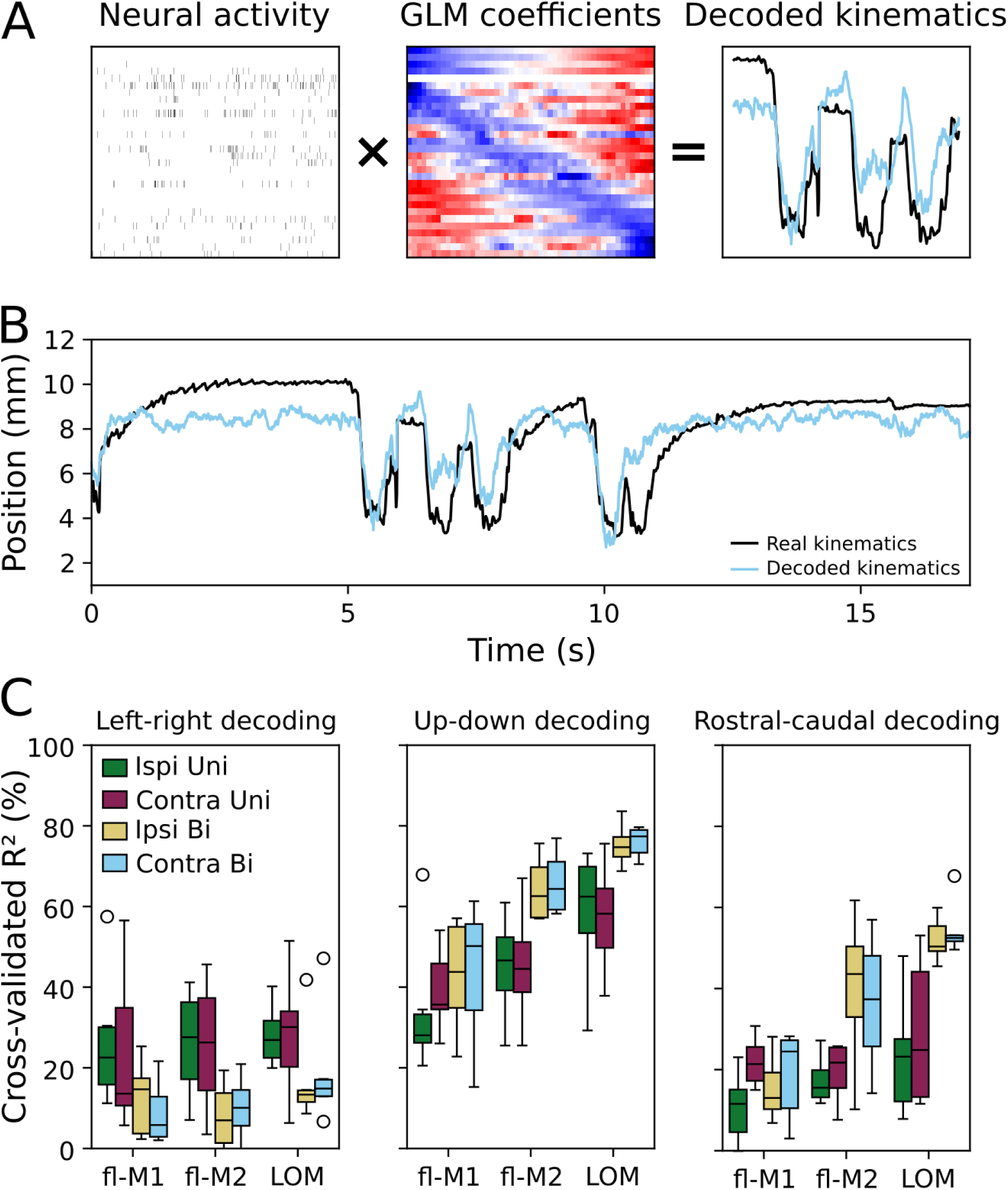
General linear model decoding of continuous food handling kinematics. (A) Schematic of GLM decoding. Sliding windows of neural activity (left) are multiplied by a matrix (time lags × units) of fitted model coefficients (middle) to reconstruct food handling kinematics on a moment-to-moment basis (right: kinematics are shown in black, model prediction in light blue). (B) Example kinematic trace (black) and decoded kinematics (light blue) from fl-M1 spiking activity. (C) Boxplots of average decoding accuracy (cross-validated *R*^2^) across areas (fl-M1: 7 recordings from 6 mice, fl-M2: 6 recordings from 4 mice; LOM: 11 recordings from 6 mice), dimensions, and conditions. Green: ipsilateral unimanual kinematics, purple: contralateral unimanual kinematics, yellow: ipsilateral bimanual kinematics, light blue: contralateral bimanual kinematics.

Having shown, as expected, that kinematics can be decoded from motor cortical activity, we next addressed the question of whether decoders trained on one limb generalize to the other (cross-body generalization, a measure of laterality invariance), and whether decoders trained on unimanual movements generalize to bimanual movements and vice-versa (uni-to-bimanual generalization, a measure of manuality invariance). For this analysis, we focused on the Y dimension as it was the best decoded from all three areas. To assess cross-body generalization, we decoded movements from activity recorded during each unimanual condition using the coefficients fit to the other unimanual condition (**Figure 10A**). To assess uni-to-bimanual generalization, we decoded each limb’s kinematics from the neural activity recorded during bimanual food handling using the coefficients fit during unimanual food handling with the same limb. We quantified generalization, or lack thereof, as the percentage change in cross-validated *R*^2^ going from the same laterality or same manuality decoder to the other laterality or manuality. A change in *R*^2^ of zero represents perfect generalization (i.e., total invariance), whereas negative values indicate a difference in how movements can be read out from neural activity between limbs/conditions. A drop to zero accuracy would give a score of −100% (i.e., total specificity).

**Figure 10:**
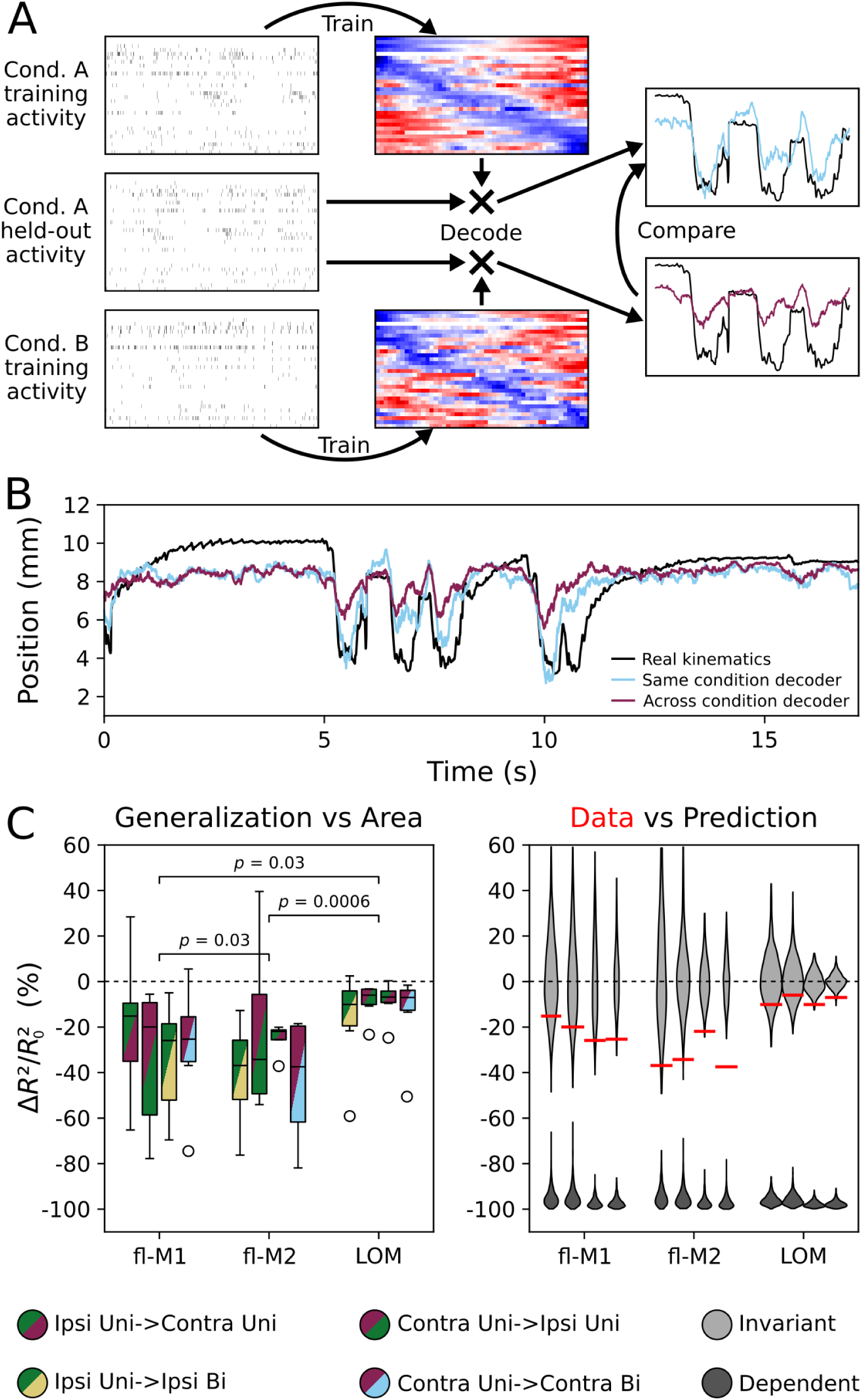
Kinematic decoding is preserved between conditions in LOM but not forelimb motor cortices. (A) Schematic of decoder generalization. Coefficients fit to neural and kinematic data from one condition (e.g. unimanual contralateral) are used to decode neural data from another condition, i.e. unimanual handling with the other hand or bimanual handling with the same hand. The cross-condition *R*^2^ is then compared to the *R*^2^ for the within-condition decoder for a held-out dataset from the same condition as the within-condition training data. (B) Example of decoder generalization. The same bimanual contralateral kinematic trace from Figure 9A is shown in black, along with its decoding from the decoder trained on bimanual contralateral data in light blue. The purple trace shows the kinematics decoded from the same activity with the decoder trained in the contralateral unimanual condition. (C) Left: average percentage change in decoding accuracy (Δ*R*²/*R*²₀) when generalizing across limbs and conditions for each area (fl-M1: 7 recordings from 6 mice, fl-M2: 6 recordings from 4 mice; LOM: 11 recordings from 6 mice). Right: expected distributions of Δ*R*²/*R*²₀ under perfect generalization (light gray) and no generalization (chance cross-condition decoding accuracy, dark gray), compared to the real data (red bars: condition pair/area medians from left panel)

**Figure 10B** shows an example of decoder generalization. The reconstruction from the across condition decoder does not follow the real kinematics as closely as that from the within-condition decoder, hence in this case there is a large percentage decrease in decoding accuracy. Across all recordings from all areas, and for all four generalizations (ipsilateral to contralateral, contralateral to ipsilateral, ipsilateral to bimanual, contralateral to bimanual), there was a negative percentage change in decoding accuracy (fl-M1: −23%, fl-M2: −35%, LOM: −7%; averages across condition pairs) compared to predictions made by within-condition decoders on held-out data (**Figure 10C**). Nevertheless, generalization was better than expected for completely dependent decoding in all areas (all *p* < 0.001). The magnitude of the decrease varied according to area (linear mixed-effects model, **Table 2**) but not between limbs or types of generalization (cross-body versus uni-to-bimanual). Comparing generalization in each area to bootstrap distributions assuming perfectly invariant decoding showed that, averaging across conditions, generalization in fl-M2 was the furthest from invariant (permutation test; fl-M1 vs fl-M2: *p* = 0.03, fl-M2 vs LOM: *p* = 0.0006; **Methods**), followed by fl-M1, with LOM the closest to invariant (fl-M1 vs LOM: *p* = 0.03; **Figure 10C**). In summary, mappings from neural activity to kinematics remain the most consistent across conditions in LOM, but change more between conditions in fl-M1 and especially fl-M2.

## DISCUSSION

We explored neural activity in three areas of motor cortex as mice use their ipsilateral, contralateral, or both hands to hold and handle food. All three areas showed large increases in activity during oromanual/ingestion, underscoring the widespread distribution of dexterity-associated activity across the large expanse of motor cortex occupying most of the frontal cortex in the mouse. In both unimanual and bimanual cases, movements of both ipsilateral and contralateral limbs could be decoded from spiking activity of multiple cortical areas. In fl-M1 and fl-M2, a majority (51-55%) of units were significantly modulated during only one of ipsilateral, contralateral, or bimanual movements. Of fl-M1/M2 units that showed significant differences in firing between conditions, the majority (63-66%) preferred contralateral and/or unimanual movements. At a population level, activity in these areas reorganized according to which hands were moving, such that decoders trained for one hand or one manuality generalized only partially to the other. Hence movement decoding displayed a mix of condition dependence and invariance. In LOM, by contrast, forelimb movement decoding was largely condition invariant. A plurality (37%) of units significantly increased in firing in all three conditions, >90% of units showed no significant preference for one limb or the other, 85% showed no significant preference for uni- or bimanual movements, and movements of both limbs independently or together could be decoded almost identically.

A limitation of our study, common to correlational studies of neural spiking activity, is that just because a variable can be decoded from spiking activity, that does not mean that is what is being encoded. As previously noted (Barrett et al., 2022), although some component of the activity in LOM may relate to ongoing jaw movements, the jaw is most active (in the form of rhythmic chewing) during the holding/chewing mode, when both the hands and LOM activity are relatively quiescent. During oromanual/ingestion epochs, when the hands and LOM are continuously active, masseter events occur as infrequent, irregularly timed bites (Barrett et al., 2024). Similarly, even if cortex is encoding hand movements, this does not imply that it is directly commanding the muscles, as such encoding may take place in an output-null subspace (Kaufman et al., 2014). Indeed, silencing of fl-M1 during food handling suggests a selective influence on forelimb movements, reflected in a reduction in regrips (Barrett et al., 2022) and change in EMG amplitude (Xing et al., 2024) rather than a complete cessation of the behavior. Such encoding may play a similar role to that proposed for rhythmic activity in forelimb motor cortex during skilled locomotion, in which it gates cortically generated adjustments to subcortically driven movements (Kirk et al., 2025). In other words, maintaining an encoding of ongoing subcortically driven movements may be necessary for motor cortex to generate the appropriate adjustment to motor output in circumstances where it is engaged (Dragoi et al., 2026; Miri et al., 2017).

Consistent with previous studies of bilateral forelimb movement related activity in rodents (Han et al., 2024; Handa et al., 2024; Jeong et al., 2021; Rios et al., 2019; Soma et al., 2017, 2019), we found a mix of laterality dependence and invariance in fl-M1 and fl-M2, with some units responding selectively during ipsilateral or contralateral unimanual movements, and others responding during both. Our study extends these results by also characterizing bilateral forelimb movement related activity during the same task performed both unimanually and bimanually, finding that activity in fl-M1 and fl-M2 also varies according to manuality. Unlike studies using unimanual lever press tasks with either forelimb in rats, which find greater contralateral bias in fl-M1 compared to fl-M2 (Rios et al., 2019; Soma et al., 2017, 2019), we largely did not find consistent differences between fl-M1 and fl-M2. This may reflect species differences, differences in task demands, or simply a lack of statistical power. Consistent with our results, in a sequential bimanual lever press task in mice (Jeong et al., 2021), activity was very similar in fl-M1 and fl-M2.

What does the lack of clear and consistent differences in laterality and manuality biases between fl-M1 and fl-M2 imply about their relation to each other? These two areas are defined as distinct areas of rodent frontal cortex that send projections to cervical spinal cord and from which forelimb movements can be evoked (Neafsey & Sievert, 1982; Nudo & Masterton, 1990; Tennant et al., 2011). Of these, fl-M2 is often considered a supplemental or premotor area by analogy with primate cortex, although the degree of homology is not clear (Barthas & Kwan, 2017). In agreement with a premotor role, fl-M2 exerts relatively more influence on fl-M1 than vice-versa during reaching (Saiki-Ishikawa et al., 2025), but this hierarchy reorganizes when task demands change, as during skilled climbing (Kristl et al., 2025). In a delayed cued push/pull task, inactivation of either area was sufficient to impair sensory discrimination and action execution, and simultaneous inactivation of both was necessary to impair maintenance of a motor plan, again at odds with a clear premotor versus primary role for one area or the other (Morandell & Huber, 2017). Thus, the similarities between fl-M1 and fl-M2 that we observed during food handling are more consistent with flexible, recurrent interactions between these two areas than with a strict premotor to primary motor hierarchy.

Laterality and manuality dependent information in primary and premotor forelimb cortical areas has also been found in primate studies (Ames & Churchland, 2019; Cross et al., 2020; Kermadi et al., 1998, 2000; Rokni et al., 2003; Willett et al., 2020; Zimnik et al., 2024). How does such condition-dependent information arise and what functional role might it serve? A trivial explanation is that, if some component of output-null processing in motor cortex is dedicated to maintaining an accurate estimate of the current and/or upcoming state of the motor plant, then different movements of different limbs would naturally evoke different firing patterns. However, maintaining separable information about what each limb is doing may facilitate bimanual coordination, and accordingly forelimb motor cortex has been implicated in bimanual control in rodents (Igarashi et al., 2019; Jeong et al., 2021) and primates (Kang et al., 2025). Similarly, interhemispheric communication has been proposed to sculpt activity in each hemisphere to facilitate task relevant movements and suppress competing outputs (Carson, 2020; Longo, 2025), but which movements should be suppressed versus permitted depends on context (i.e., unimanual versus bimanual), and hence maintaining separable information about manuality is also necessary.

Laterality dependence of forelimb movement related activity in LOM has not previously been investigated, but activity relating to tongue movements in LOM (Xu et al., 2022) and adjacent anterolateral motor cortex (ALM) (Guo, Li, et al., 2014; Li et al., 2015) has been studied. In these areas, in contrast to the largely laterality-invariant forelimb movement activity that we found, neural activity showed strong selectivity for left versus right licks (Li et al., 2015; Xu et al., 2022), although there is also a large non-selective, movement-related (‘go’) signal (Inagaki et al., 2022). Interpreted somatotopically, this difference is consistent with LOM being considered a primary motor cortex area specifically for the tongue and jaw, and not the forelimb. However, recent evidence strongly implicates LOM in coordination of hand and orofacial movements (Barrett et al., 2022; Huang et al., 2025; Khanal et al., 2024; Yang et al., 2023). Considering the relevance of laterality and manuality dependent information to different types of coordination offers an alternative interpretation, which our findings favor. In directional licking, the body side to which movements are directed is integral to task success. In oromanually coordinated food handling, in contrast, the key task variable is proximity of the hand(s) to the mouth; the particular direction the food is coming from (laterality) and the number of hands involved (manuality) are less relevant. The high degree of laterality and manuality invariance observed in LOM for forelimb movements is thus entirely expected under a framework in which activity in LOM encodes relevant movement variables, regardless of effector, for both skilled licking and food handling. The tonically elevated activity in LOM throughout oromanual/ingestion epochs is also consistent with this hypothesis, i.e., LOM remains active as long as the hands are at the mouth (Barrett et al., 2022).

In summary, we have shown that activity in fl-M1 and fl-M2 varies according to which limbs are moving as mice dexterously handle food with one hand, the other, or both. At the same time, activity in LOM tracks how close the hands are to the face at any given moment, invariant to laterality and manuality. These distinct patterns of laterality and manuality dependence and invariance are consistent with distinct roles for each of these areas in different types of coordination – bimanual and oromanual – during natural behavior. Future work disrupting inter-area and interhemispheric communication during coordinated behavior will be necessary to confirm these roles, and to elucidate the circuit and computational mechanisms by which dependent versus invariant movement activity is constructed and maintained.

## MATERIALS AND METHODS

### Subjects

This study used experimentally naïve mice of both sexes on a C57BL/6 background (stock no. 000664, The Jackson Laboratory, Bar Harbor, Maine; Table 1) aged 109-231 days postnatal and weighing 18.3-28.4 g at the time of recording (21.7-33.5 g before food restriction, minimum 56 days postnatal and 20 g at the time of surgery). Mice were selected for weight as larger mice were found to have better surgical outcomes. Mice were bred in-house, housed in groups with a 12-hour reverse light/dark cycle, and had free access to food and water prior to food restriction (see below). As many brain areas as possible (of the total of six bilateral representations of the three areas of interest) were recorded from each mouse, hence no randomization to cohorts was necessary and mice were used as they became available. Experiments were conducted during the dark phase of the mice’s light cycle. Studies were approved by the Northwestern University IACUC and complied with the animal welfare guidelines of the National Institutes of Health and Society for Neuroscience.

### Surgical procedures

Under deep isoflurane anesthesia (3% induction, 0.8-1.5% maintenance), the skin over the cranium and muscles to be implanted was shaved and cleaned, mice were placed in a stereotaxic frame (Model 900, David Kopf Instruments, Tujunga, CA), and a midline incision was made to expose the cranium. The periosteum was removed, the skull gently scraped to improve adhesion, and a titanium head-fixation bar (0.875 × 0.187 inches, cut by water-jet from 0.08 inch Ti-6Al-4V sheet, Big Blue Saw, Atlanta, GA) was placed on top of lambda, perpendicular to the central suture, and affixed using dental cement (C&B Metabond, Parkell, Edgewood, NY).

The cranial incision was then sutured to close the wound margins and cover any exposed cranium or muscle not covered by dental cement and/or the head-bar. Mice were given 1 mg/kg buprenorphine intraperitoneally and a small amount of bupivacaine (<2 mg/kg) under each incision site preoperatively and 20 mg/kg meloxicam subcutaneously postoperatively as analgesia along with 10 mg/kg enrofloxacin subcutaneously, followed by a second and third doses of meloxicam and enrofloxacin 24 and 48 hours later. Mice were single-housed following head-bar mounting.

One day prior to first recording, mice were deeply anaesthetized with a cocktail of 80–100 mg/kg ketamine and 5–15 mg/kg xylazine injected intraperitoneally. A craniotomy or craniotomies were opened over the area(s) to be recorded using a dental drill (EXL-M40, Osada, Los Angeles, CA). The cortical surface was covered with Kwik-Sil (World Precision Instruments, Sarasota, FL) and mice were allowed to recover.

### Electrophysiological recordings and analysis

The linear arrays used were either 32-channel silicon probes with ∼1 MΩ impedance and 50 μm spacing in linear configuration (model A1×32-15mm-50-177-A32, NeuroNexus, Ann Arbor, MI) or 64-channel probes in two staggered columns, each with 46 μm vertical spacing, and 23 μm horizontal spacing between columns (A1×64-Poly2-6mm-23s-160-A64). Each probe was mounted on a linear translator (MTSA1, ThorLabs) that was in turn mounted on a 3-axis manipulator (MP285, Sutter, Novato, CA). Probes were positioned at the recording sites stereotactically based on the stereoscopically visualized location of bregma using the three axes of the Sutter manipulator, then slowly inserted into the cortex using the linear translator at a rate of 2 μm/s (controlled by software) to a nominal depth of 2000 μm from the pia. Target coordinates for the three areas were as follows: fl-M1, 0.2 mm anterior-posterior (AP), 1.3 mm medial-lateral (ML); fl-M2, 2.0 mm AP, 1.2 ML; LOM, 1.8 AP, 2.5 ML. For LOM recordings, the probes were tilted by ∼30° off the vertical axis in the coronal plane for alignment with the radial axis of the cortex. For fl-M1 and fl-M2 the probes were inserted perpendicularly to the horizontal plane (for unilateral recordings of either area with LOM) or ∼15° off the vertical axis in the coronal plane (for bilateral recordings, to avoid headstage collision). For simultaneous unilateral recordings for fl-M1 and fl-M2, the fl-M1 probe was inserted vertically and the fl-M2 probe was tilted by ∼15° off the vertical axis in the sagittal plane. At the end of each experiment, the probes were removed, the craniotomy re-sealed with Kwik-Sil, and the mouse returned to its home cage. A maximum of three recording sessions were performed per mouse, on subsequent days.

Signals were amplified and hardware bandpass-filtered (2.5 Hz to 7.6 KHz) using RHD2132 headstages (Intan Technologies, Los Angeles, CA) and acquired at 30 kHz using an RHD2000 USB Interface Evaluation Board (Intan). Data was recorded using the Intan experimental interface evaluation software, triggered by the same trigger used to control video recording. To synchronize videos to electrophysiological recordings, the frame sync signal from the camera was recorded as a digital input to the RHD2000. RHD files recorded by the Intan software were concatenated using spikeinterface (Buccino et al., 2020) and spike sorted using Kilosort 4 (Pachitariu et al., 2024). Results from Kilosort were manually verified using phy (https://github.com/cortex-lab/phy) as follows. Units with waveforms spanning more than 3 adjacent channels or with atypical waveform shapes were rejected as artifactual. Units displaying a clear refractory period (<1% of spikes within 1 ms) were classified as single units. All other units were classified as multiunit activity. Multiunits on the same channel with similar waveform shapes were merged. Single units were merged only if they were on the same channel, displayed similar shapes, and had refractory crosscorrelograms. Single units and multiunits were included in all analyses presented. Only data from probe recordings from which at least 10 units could be isolated were included.

### Behavioral training and videography

Mice were acclimatized to head-fixation as previously described (Barrett et al., 2022). Briefly, following at least one week of recovery from head bar implantation, mice were placed on food restriction, receiving a limited amount of standard rodent diet each day to maintain their bodyweights at 85-90% of pre-restriction bodyweights. Mice were monitored throughout the food restriction period for signs of ill-health (Guo, Hires, et al., 2014), and body condition scores (Ullman-Culleré & Foltz, 1999) were taken each day. No signs of ill-health were observed, and no mouse fell below a body condition score of 3 throughout the study. Once on food restriction, mice were familiarized with the apparatus and handling by the experimenter, then exposed to progressively longer durations of head-fixation until they became comfortable handling and consuming food items presented to them while head-fixed. Mice were fed shelled sunflower seed kernels (S5137-1, Bio-Serv, Flemington, NJ) or 45 mg grain-based dustless precision pellets (F0165, Bio-Serv). While head-fixed, mice were enclosed in a custom 3D printed hut, as previously (Barrett et al., 2022, 2024). To encourage unimanual food handling, the cylindrical hand rest incorporated into the hut was replaced with a pair of 3D printed hand blockers that could rotate back and forth and slide from side to side (**Figure 2A**). The hand blockers were designed with multiple protruding flaps that, when rotated into place, prevented access by one hand to the area immediately in front of, below, and to the side of the face.

Videos were acquired with a high-speed CMOS-based monochrome video camera (Phantom VEO 710L, Vision Research, Wayne, NJ) at 1000 frames per second (fps), 999.6 µs exposure time, and 1024 × 512 pixel field of view. Two oblique views of the mouse were obtained by mounting two 50 × 50 mm flat enhanced aluminum surface mirrors (#43-876, Edmund Optics, Barrington, NJ) and a 50 mm anti-reflection coated equilateral prism (#49-435, Edmund Optics) in the camera optical path. A prime lens (Nikon AF Micro-NIKKOR 60mm f/2.8D, Nikon, Tokyo, Japan) was mounted on the camera body. The mouse was illuminated from both sides and slightly below using two red LEDs (M660L1 and MLEDC25, ThorLabs, Newton, NJ). Camera and video recording settings were controlled with Phantom Camera Control Application v3.5 (PCC, Vision Research). Video was recorded to the camera memory and then saved to disk as uncompressed Phantom Cine files, later converted to H.264-encoded MP4 files. For bimanual food handling, video recording was manually triggered by a TTL pulse delivered from an NI USB-6229 data acquisition board (National Instruments, Austin, TX) once the mouse had successfully retrieved a food item in both hands and begun eating. For unimanual food handling, video was triggered on the first successful food acquisition and recording continued over multiple food presentations until the camera memory was full. In both cases a one second video pre-trigger buffer was used.

Videos were cropped to isolate each view using ffmpeg (ffmpeg.org). Markerless tracking of the nose, the tip of the lower jaw, and the second through fourth digits (D2-4) on each hand was performed using DeepLabCut (Mathis et al., 2018; Nath et al., 2019). From these two sets of 2D trajectories for each body part, 3D trajectories were reconstructed using Anipose (Karashchuk et al., 2021). Anipose’s camera model was calibrated for each experimental session using videos of a ChAruCo board at various angles captured at the end of the session without adjusting the lens settings. As previously (Barrett et al., 2020, 2022, 2024) we focused our analysis on *D_hand-hand_*, the 3D Euclidean distance between the third digits (D3) of each hand, and *D_both-nose_*, the distance between the mid-point of the two D3s and the nose. The distance between D3 and the nose was also calculated separately for the left and right hands as *D_left-nose_* and *D_right-nose_*, respectively.

### Histology

Following the final recording from each mouse, the mouse was sacrificed by ketamine/xylazine overdose followed by transcardial perfusion following standard procedures. The brain was then quickly dissected and stored overnight in 4% paraformaldehyde in phosphate-buffered saline (PBS), before being washed and stored in PBS with 0.02% w/v sodium azide the next day. Post-hoc localization of recording site locations was then performed as previously (Barrett et al., 2022). Briefly, fixed brains were sliced coronally at 100-150 µm thickness and imaged under brightfield and fluorescent illumination to identify DiO, DiI, or DiD (Vybrant Multicolor Cell Labeling Kit, Invitrogen, Carlsbad, CA) fluorescence left behind by the dye-coated probes. Recordings with an AP coordinate less than 0.5 mm anterior and an ML coordinate less than 1.7 mm lateral were assigned to fl-M1 (Barrett et al., 2022). Recordings with an AP coordinate greater than 1.5 mm anterior were assigned to fl-M2 if their ML coordinate was less than 1.5 mm lateral and LOM otherwise (Yang et al., 2023). Recordings outside these boundaries were not analyzed. The median recording locations were as follows – fl-M1: 0.0 AP, 1.4 ML; fl-M2: 2.0 AP, 1.1 ML; LOM: 1.8 AP, 2.3 ML (**Figure 4—figure supplement 1**).

### Ethogramming

Semi-automated ethogramming of hand kinematics into holding and oromanual modes was performed as previously (Barrett et al., 2022, 2024). Briefly, videos were segmented into holding/chewing, oromanual/ingestion, or other based on manual thresholding of *D_left-nose_* and *D_right-nose_*, then characteristic model functions (sigmoid for transport-to-mouth, exponential for lowering-from-mouth) were fit to each transition to more precisely determine their start and end. Whereas previously we excluded unimanual handling by including it in the ‘other’ category, here we constructed a three-state ethogram for each hand separately and combined these into a seven state ethogram (left holding/chewing, right other; left oromanual/ingestion, right other; right holding/chewing, left other; right oromanual/ingestion, left other; both holding/chewing; both oromanual/ingestion; both other). Transitions where one or both hands were rising (from other into holding/chewing or oromanual/ingestion, or from holding/chewing into oromanual/ingestion) were fit with sigmoid functions to the hand-nose distance of the upwardly moving hand(s) and all other transitions were fit with exponential functions to the hand-nose distance of the downward-moving hand(s).

### Analysis of spiking activity

Only recordings with at least ten units and at least five transports-to-mouth of each handedness were analyzed. For analysis of spiking activity associated with transports-to-mouth, peri-event time histograms (PETHs) were constructed for each unit by aligning spike timestamps to the onset of each transport-to-mouth and binning the aligned spike times in 20 ms bins from one second before to one second after the transport-to-mouth onset. Separate PETHs were constructed for left, right, and bimanual transports-to-mouth. Only transitions from holding/chewing to oromanual/ingestion with the same hand(s) were considered, e.g. left holding/chewing to left oromanual/ingestion. Uni- to bimanual transports-to-mouth, hand swaps, and transitions from hands resting on the armbar into oromanual/ingestion were excluded from analysis.

Significance of PETHs was assessed as follows. For each handedness of transport-to-mouth, sham event times were generated uniformly distributed throughout each recording in equal number to real transports-to-mouth of that handedness. This process was repeated 1000 times and, for each iteration, the maximum of the PETH in the range from 500 ms before to 500 ms after transport-to-mouth onset was calculated to generate a bootstrap distribution of maximum mean firing rates for each unit. This distribution was used to assign a right-handed *p*-value to each unit’s PETH equal to the number of bootstrap maximum mean firing rates greater than that unit’s actual maximum mean firing rate from −500 ms to +500 ms, divided by 1000. PETHs with a *p*-value less than 0.001 were considered significant. Units with a significant maximum mean firing rate for any condition were considered significantly responsive. Only recordings with at least five significantly responsive units were used in subsequent analysis of PETHs.

To assess laterality/manuality preferences, a preference index was calculated for each unit as follows. First, all PETHs were baseline subtracted on a per-event basis by subtracting the mean firing rate from 1 second to 200 ms before the corresponding transport-to-mouth, excluding any preceding oromanual/ingestion epochs. Then, the mean PETH was taken over all events recorded on the same day and the time of maximum mean firing over the interval −200 ms to +500 ms found. Each unit’s response to a given handedness of transport-to-mouth was defined as the area under the curve of the mean PETH in a 100 ms (i.e. 5 bin) window surrounding that unit’s time of maximum mean firing. A preference index (PI) was calculated for each unit as (*A*-*B*)/(*A*+*B*), where *A* and *B* are that unit’s responses in the conditions being compared. Cumulative distributions of preference indices were calculated per recording, then the mean was taken over probes, days, and mice. Average cumulative distribution functions were compared using the two-sample Kolmogorov-Smirnov distance (maximum absolute difference between the two distributions) and significance was assessed using a permutation test, randomly reassigning the area label for each unit and repeating for 10000 iterations.

Units were assigned as having a strong laterality/manuality preference if the *p*-value of a Mann-Whitney *U* test comparing their responses in a given condition pair was less than 0.05. The *p*-values used for this classification were not corrected for multiple comparisons as permutation testing showed that doing so introduced more false negatives than it excluded false positives. Two-dimensional histograms of preference categories were calculated per recording, averaged over probes, days, and mice, and compared using the earth mover’s distance (EMD) (Peleg et al., 1989). The metric used to optimize the EMD was the Euclidean distance between bins arranged as in **Figure 6F**, assigning unity distance to horizontally or vertically adjacent bins. The EMD was calculated using pyemd (Flamary et al., 2021) and significance was assessed using a permutation test, randomly reassigning the area label for each unit and repeating for 10000 iterations.

### Principal components analysis

For principal components analysis, all units’ trial-mean PETHs for a given recording were soft-normalized by dividing by their maximum firing rate plus 5 (Elsayed et al., 2016), then concatenated into a matrix **X** of dimension *T*×*N*, where *T* is the number of time points and *N* the number of units. **X** was decomposed via singular value decomposition into two rank *P* matrices **PQ** of dimension *T*×*P* and *P*×*N*, respectively. **Q** gives the projection from neural activity space into principal component space. The first principal angle between the subspaces spanned by the top principal components was calculated using the subspace_angle function in scipy. Denoting the covariance matrix of the population activity in condition A as **C_condA_** = cov(**X_condA_**,**X_condA_**), the participation ratio (Gao et al., 2017) was calculated as [tr(**C_condA_**)]^2^/tr(**C_condA_**^2^). Denoting the *i*th row of **Q** in condition A as **q***^i^*_condA_ the *i*th singular value of **C_condA_** as *σ^i^*_condA_, the between condition alignment index for the top 10 principal components in conditions A and B was calculated as the variance explained by the activity from condition A projected onto the top 10 principal components from condition B, divided by the variance explained by the activity from condition A projected onto the top 10 principal components from condition A; i.e., **Σq***^i^***_condB_C_condA_q***^i^***_condB_**^T^|/**Σ|***σ^i^*_condA_| (Elsayed et al., 2016). The alignment index approaches unity as the number of components used to calculate it approaches the maximum dimensionality of the raw data, hence only recordings with at least 40 units were included in analyses related to principal components. To increase the amount of available data, we included recording sessions where only a subset of the three conditions were successfully captured, but increased the required number of transports-to-mouth for inclusion to 15. For aggregating data across recordings, conditions with less than this number transports-to-mouth were excluded from the aggregation (i.e., treated as NaN).

In the finite data regime, subspace angles of 0°/90° and alignment indices of 1/0 are unlikely even for samples from perfectly aligned/orthogonal subspaces. Hence, to estimate expected distributions of subspace angle and alignment index under these assumptions, we generated bootstrap distributions as follows. For each recording, to simulate invariant encoding, we split the transport-to-mouth aligned activity from two conditions of interest into two halves, calculated a weighted average (inversely weighted by the number of trials so both conditions contribute equally), performed principal components analysis on the resulting average PETHs, and calculated the subspace angle and alignment index for the two halves. To simulate orthogonal subspaces for ipsilateral and contralateral data (laterality dependence), we again split the data in halves, using one half to find two separate orthogonal projections that respectively maximize the explained variance of the ipsilateral and contralateral data (similar to the procedure used to orthogonalize the preparatory and movement subspaces in Elsayed et al., 2016). Projections were found via Riemannian manifold optimization using pymanopt (Townsend et al., 2016). Then we projected the remaining half of the data for each condition into these subspaces and performed the subspace angle and alignment index analyses on these projected data. Manuality dependence was simulated similarly by finding two orthogonal projections: one that maximizes the sum of the explained variance for the ipsilateral and contralateral data, and one that maximizes the explained variance for the bimanual data. Both of these were repeated 1000 times for each recording, and the resulting subspace angles and alignment indices averaged as for the real data (over probes, days, and mice) to produce the expected distributions shown in **Figure 7G-H**. To assess whether observed subspace angles or alignment indices better matched a given prediction, we calculated the difference between the real subspace angle/alignment index and the median predicted subspace angle/alignment index, averaged these differences over probes, dates, recordings, mice, and conditions, then took the difference between the differences. Finally, this was compared to a bootstrap distribution computed by performing the same calculation with permuted area labels, for 10000 iterations.

### Analysis of population correlation structure and decoding of forelimb kinematics

For the population correlation structure and GLM decoding analysis, spike trains were binned into 5 ms bins and kinematics (X, Y, and Z positions of D3 of each hand) were downsampled to 200 frames per second, then both sets of data were separated into epochs corresponding to ipsilateral food handling (i.e. ipsilateral holding/chewing and ipsilateral oromanual/ingestion), contralateral food handling, and bimanual food handling. Sections corresponding to non-food handling behavior (e.g. resting hands on the arm bar or postural adjustments) were not included in this analysis. Units with an all time average firing rate less than 0.5 Hz were excluded from these analyses. After applying these exclusion criteria, all recordings with at least 10 units and where all three conditions had at least 5 transports to mouth and 1000 time bins were used for correlation and decoding analyses.

For analyzing population correlation structure, binned spike trains for each unit in each condition were concatenated into a single long vector. Then, for each condition, Pearson’s product-moment correlation coefficient (*ρ*) was calculated between all units’ firing vectors. The upper triangle of the resulting matrix was then reshaped into a vector defining the population correlation structure for each condition. Finally, *ρ* was calculated between the population correlation structure vectors for each pair of conditions to assess the between-condition similarity.

For decoding, as previously, we used ridge regularized GLMs on lagged windows of neural activity to decode kinematics (Barrett et al., 2022). The decoding window spanned 200 ms before to 50 ms after the kinematic time bin being decoded. The datasets for each of the three conditions were subdivided equally into non-overlapping hyperparameter tuning, training, and testing sets. Separate GLMs were fit to each kinematic trace in each condition, except that GLMs were not fit to ipsilateral kinematics during contralateral unimanual food handling and vice versa due to the frequent occlusions of the unused hand. The optimal ridge parameter *λ* for each GLM was found by grid search with five-fold cross-validation over the hyperparameter tuning set. Then, GLMs were re-fit using the chosen *λ* to the training set. Finally, decoding performance was assessed by calculating the coefficient of determination (*R*^2^) between the real kinematics from the testing set and the decoded kinematics using the corresponding neural data from the testing set (cross-validated *R*^2^).

To assess cross-body generalization, we decoded kinematics from neural activity from one limb’s unimanual testing set with the GLM fit to the other limb’s unimanual training set. To assess uni-to-bimanual generalization, we decoded kinematics from neural activity from one limb’s bimanual testing set with the GLM fit to the same limb’s unimanual training set. In both cases, generalization was calculated as the percent change in *R*^2^ going from the same limb/condition decoder to the other limb/condition decoder (Δ*R*²/*R*²₀).

To estimate distributions of Δ*R*²/*R*²₀ under the assumptions of perfect generalization (total invariance) or no generalization whatsoever (total specificity, or chance decoding), we generated bootstrap distributions as follows. For each recording, we took the testing set from a given condition and split it randomly in half (by chunking the data into 20^th^s, permuting them, and concatenating the first 10 and last 10 permuted chunks). To simulate perfect generalization, we decoded each half’s kinematics from the same half’s neural data and calculated Δ*R*²/*R*²₀, arbitrarily choosing one half as the within-condition set and the other as the cross-condition set. To simulate no generalization, we decoded the first half’s kinematics from the second half’s neural data, and compared this to decoding from the first half’s neural data. For both simulations, the same decoder was used, i.e. the one trained on the same condition the testing set data was taken from. Each simulation was performed 1000 times per recording, and the resulting Δ*R*²/*R*²₀ values averaged over probes, days, and mice to obtain the final bootstrap distributions shown in **Figure 10C**. To assess whether generalization in one area was further from or closer to the invariant prediction than another area, we calculated the difference between the real generalization and the median predicted generalization assuming invariance, averaged these differences over probes, dates, recordings, mice, and conditions, then took the difference between the differences. Finally, this was compared to a bootstrap distribution computed by performing the same calculation with permuted area labels, for 10000 iterations.

### Quantification and statistical analysis

Time series data are presented in figures as mean ± S.D. except where noted and scalar data is presented as median ± interquartile range (IQR) boxplots with whiskers out to the farthest datapoint within 1.5*IQR of the nearest quartile, and all points outside this range shown as outliers. In general, averages were taken first over trials, then over units recorded on the same probe, then over probes recorded on the same day for bilateral recordings of the same area (reversing the ipsi-and contralateral conditions as appropriate), then over recordings of the same over separate days from the same mice, then finally over mice. Not all areas were successfully recorded the same number of times from all mice, hence the dataset analyzed in this paper represents an unbalanced, incomplete block design. Thus most statistical tests reported in this paper used linear mixed-effects models, where the comparisons of interest (e.g. between areas, between limbs, between conditions) were included as fixed effects and individual mice were included as random effects. Main effects were included for all models, and interaction terms were only included for two-way models. For follow-up pairwise testing, we also used linear mixed-effects models (except where noted), including only the two levels of the factor of interest being compared, and ignoring the other factors. For these tests, *p*-values were corrected using the Holm-Bonferroni method.

## ACKNOWLEDGEMENTS

We thank Daniela Piña Novo for helpful comments, as well as Miraya Baid, Megan Martin, and Rita Fischer for technical assistance. Funding was provided by the National Institute for Neurological Disorders and Stroke (grant numbers 5R37NS061963-17 and 1R21NS135642-01).

## AUTHOR CONTRIBUTIONS

Conceptualization: JB

Data curation: JB

Formal analysis: JB

Funding acquisition: GS

Investigation: JB

Methodology: JB, JG, AM

Project administration: JB, GS

Resources: GS

Software: JB

Visualization: JB

Writing – original draft: JB

Writing – review & editing: JB, JG, AM, GS

## DATA AND CODE AVAILABILITY

Data and code necessary to reproduce the results presented in the figures will be made available at Zenodo prior to publication.

---

**Figure 4—figure supplement 1:**
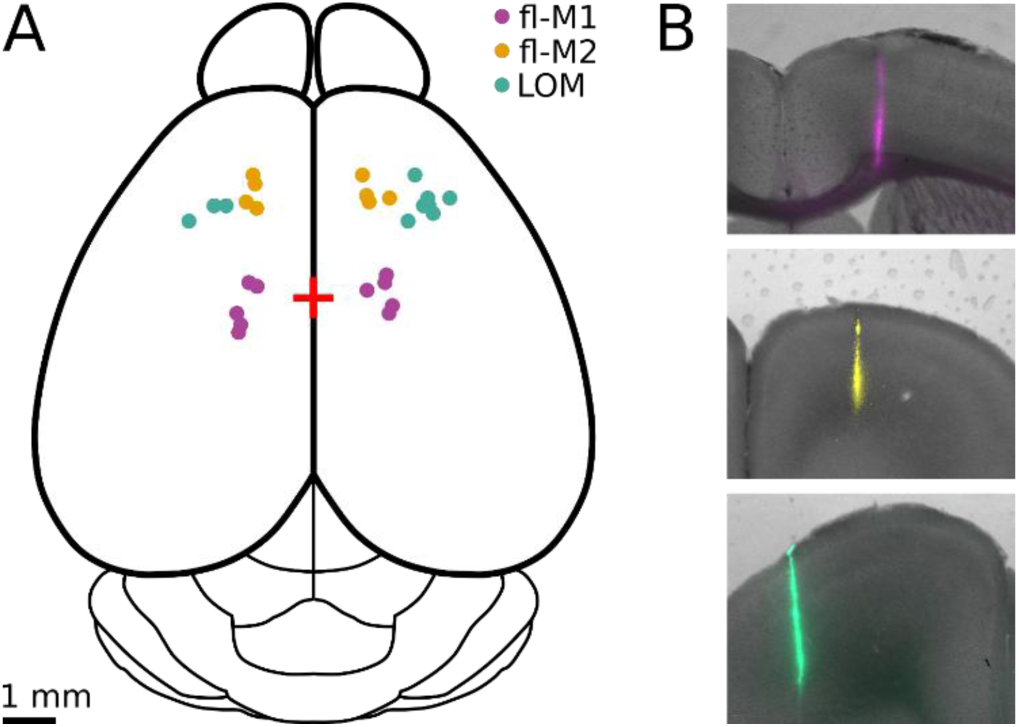
recording site locations. (A) Schematic dorsal view of a mouse brain annotated with locations of all fl-M1 (violet), fl-M2 (gold), and LOM (teal) recordings analyzed in this paper. (B) Merged brightfield-epifluorescence images of brain slices with pseudocolored fluorescent probe track locations for example fl-M1 (top, magenta), fl-M2 (middle, yellow), and LOM (bottom, teal) recordings.

**Figure 7—figure supplement 1:**
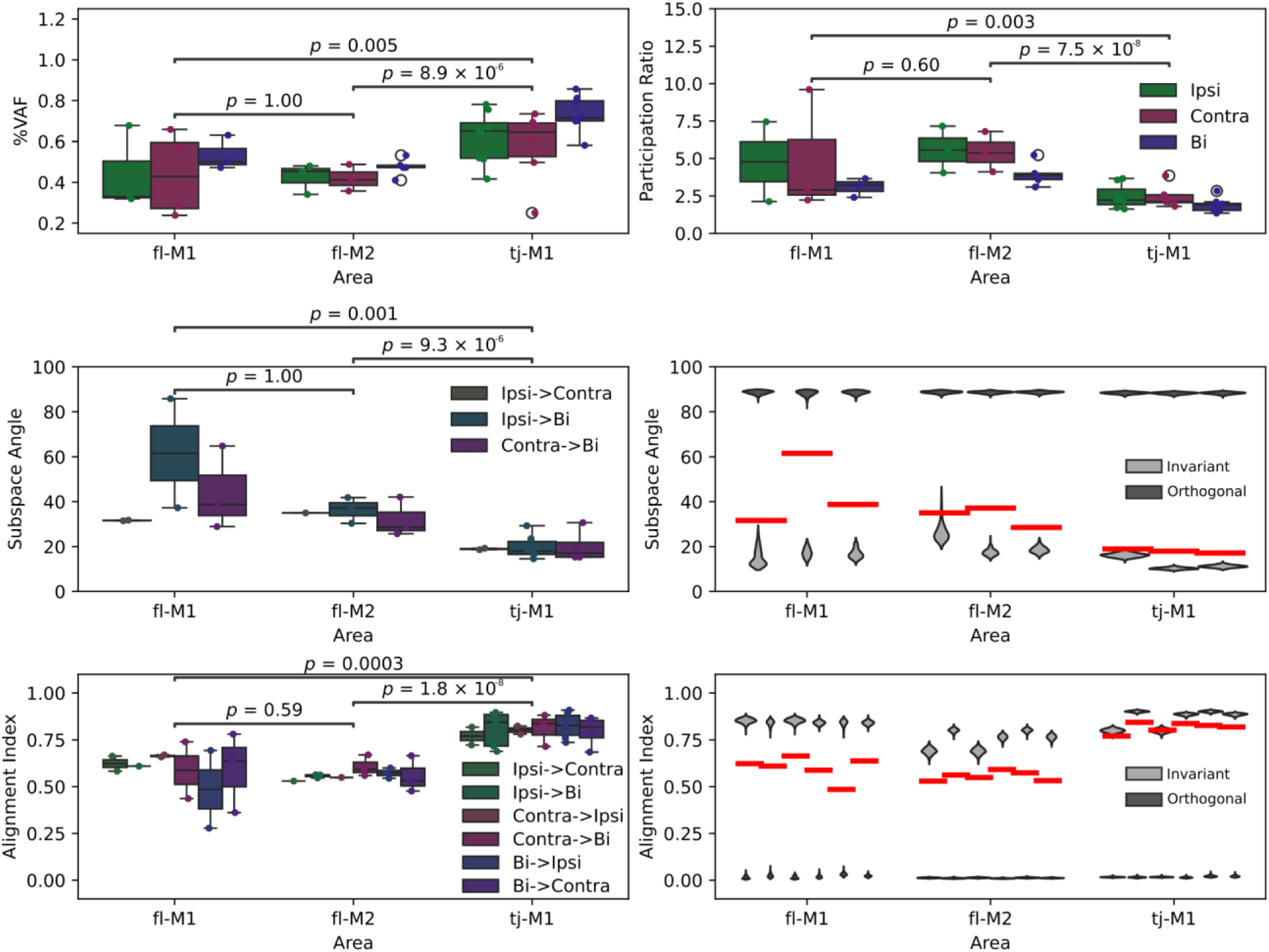
principal components analysis and related analyses without pooling fl-M1 and fl-M2. (C) Boxplots of fraction of explained variance accounted for by the top principal component of transport-to-mouth aligned average activity, organized by area then condition. (D) As (C), for participation ratio. (G) Left: Boxplots of subspace angles between the top principal components for each area and condition pair. Right: Expected distributions of subspace angles under laterality/manuality invariance (light gray) and maximum laterality/manuality dependence (i.e. orthogonality, dark gray) compared to the real data (red bars: condition pair/area medians from left panel). (H) As (G) for alignment index between top ten principal components.

